# The chemical basis of metabolic interdependence in microbial communities

**DOI:** 10.1101/2022.03.14.484247

**Authors:** Akshit Goyal, Sandeep Krishna

## Abstract

Microbial communities play a crucial role in determining the dynamics of soil and marine ecosystems. They strongly influence the physiological functioning of plants and animals, for instance, nutrient uptake, stress tolerance, immune responses in the gut, lung, skin, etc. The diverse species in such communities interact both competitively as well as cooperatively. Cross-feeding, the exchange of metabolites between a pair of microbial species for mutual benefit is a common interaction that probably explains why 99% of natural bacterial species are unculturable on their own in the laboratory. Here, we provide a theoretical, network-level understanding of the conditions under which cross-feeding between a pair of microbial species can be beneficial to both. Using the known microbial repertoire of metabolic reactions, as represented in the KEGG database, we construct a large ensemble of metabolic networks designed to synthesize a set of biomass precursors from specified nutrients. We construct both autonomous networks, that can perform this task on their own, as well as pairs of cross-feeding networks that can only perform this task together but not alone. Surprisingly, we find that there exist cross-feeding pairs that produce higher biomass or energy yields than even the best autonomous networks. We show that such “outperforming” cross-feeding pairs exist only because of certain nonlinearities in the way metabolic flux is distributed in these networks. By analyzing patterns in our ensemble of networks, we propose a set of necessary and (almost) sufficient conditions that the metabolic networks have to satisfy for cross-feeding to be beneficial. These conditions are based partly on the structure of the networks and partly on the chemical and thermodynamic properties of the underlying chemical reactions, phenomenologically quantified in terms of the effect of donating or accepting metabolites on the yield of our constructed networks. Our analysis not only provides a mechanistic understanding of why cross-feeding is prevalent in microbial communities, but also provides a theoretical basis for understanding the benefit of compartmentalization of chemical reactions in a variety of contexts, for instance with mitochondrial vs. cytoplasmic metabolism in eukaryotic cells, or multi-enzyme cascade reactions in industrial contexts.

## Introduction

Naturally occurring microbial communities are genetically and metabolically diverse. A common argument to explain the prevalence of such diversity is that co-occurring microbial species can often benefit from each other through division of labour [1, 2, 3, 4, 5]. In many cases, this labour is biochemical in nature. For instance, species can benefit by *cross-feeding*, where one species releases metabolites into its environment which another can use as a nutrient [6, 7, 8]. A mutualistic interaction, where a pair of species cross-feed off each other, can offer a fitness benefit. That is, the growth rates of both species can increase due to cross-feeding when compared with autonomous culture, even when the interaction is obligate [9, 10].

Some reports suggest that the dominant reason why only about 1% of naturally occurring microbes can be cultured in the lab [11, 12] is that obligately dependent cross-feeding is prevalent in natural communities [13, 14, 15]. Others, however, argue that competitive interactions are more common between microbes in nature [16, 17] and an obligate cross-feeding interaction, albeit beneficial, would likely be unstable against environmental and genetic changes [18, 19, 20]. This is a major on-going debate.

Previous studies that have attempted to understand metabolic interdependence between microbes fall into three broad categories: (i) measuring fitness benefits in genetically engineered bacteria which survive by exchanging metabolites [9, 21, 22, 23], (ii) inferring interactions in naturally occurring communities through shotgun metagenomic sequencing [24, 25, 26, 27], and (iii) using theoretical models to understand the requirements for beneficial cross-feeding [18, 28, 29, 30, 31, 32]. These studies provide evidence both for and against: some suggest that obligate cross-feeding can help interacting species [9, 22, 23, 33], while others claim that this is likely not the case [17, 19, 20, 27]. However, these studies have some limitations: the experiments either lack a mechanistic understanding of the interaction (which metabolites are exchanged and how that benefits fitness), or are constrained by the lack of metabolic annotations in sequenced community samples; and the theoretical models rely on metabolic trade-offs that have not been experimentally verified to show the aforementioned benefits.

Here, we construct an ensemble of metabolic networks randomly assembled from the entire repertoire of prokaryotic metabolic reactions (as represented by the KEGG database). The random assembly of these networks is subject only to constraints of viability, defined as the ability to synthesize a specified set of biomass precursors from a given set of nutrients. We construct such ensembles both for autonomous networks that are viable on their own, and cross-feeding pairs of networks that are viable together but not individually. We use these ensembles to provide baseline expectations for the structure, productivity and stability of autonomous and cross-feeding pairs of metabolisms based upon the chemical reactions that comprise these networks and their stoichiometry. Strikingly, we demonstrate that some cross-feeding pairs can produce higher biomass yields than even the best autonomous networks. We identify the chemical basis for this phenomenon; that is, we provide necessary and sufficient conditions that the network pairs must satisfy for cross-feeding to be beneficial.

## Results

### Algorithmic generation reveals on the order of 10,000 unique autonomous metabolic networks viable in a given nutrient environment

We first studied autonomous networks, which can individually convert nutrients into biomass precursors and energy. We assumed a minimal nutrient medium with acetate as a carbon source and ammonia as a nitrogen source. Further, we assumed that the following “currency” molecules were always present in saturating amounts in the medium: water, carbon dioxide, oxygen, hydrogen, ATP, ADP, AMP, NAD(P)H and formate. We verified that changing the carbon and nitrogen source to glucose and glutamate, respectively, does not qualitatively affect our results (supplementary figure 4).

We then stipulate that a “viable” metabolic network must satisfy the following stringent condition: starting from only the nutrient and currency molecules, it must be able to produce *all* of a specified set of biomass precursor molecules (such as pyruvate, L-Serine, and D-Ribose-5-phosphate; see table 2 for full list of precursors we use). We have also verified that an alternate choice for the precursors does not qualitatively affect our results.

For the medium composition described above, we were able to generate 19,543 unique randomly assembled networks that satisfied this criterion. The algorithm for doing this is depicted schematically in Fig. 1 and described in detail in Methods. We note here only that: (i) the reactions comprising these networks are chosen from the entire repertoire of prokaryotic metabolic reactions in the KEGG database (henceforth referred to as the “KEGG universal chemistry”), and (ii) this algorithm generates “minimal” networks, in the sense that removal of any single reaction from each network causes it to become unviable. The latter choice was made simply to allow a more robust comparison between such autonomous networks and the cross-feeding pairs of networks we discuss in the next section. Figure 1A shows one example of such a constructed network, specifically the smallest autonomous network we generated (the one with the least number of reactions). It is interesting that so many solutions are possible within the KEGG universal chemistry for the metabolic task of converting the given nutrients into the specified biomass precursors.

**Figure 1:**
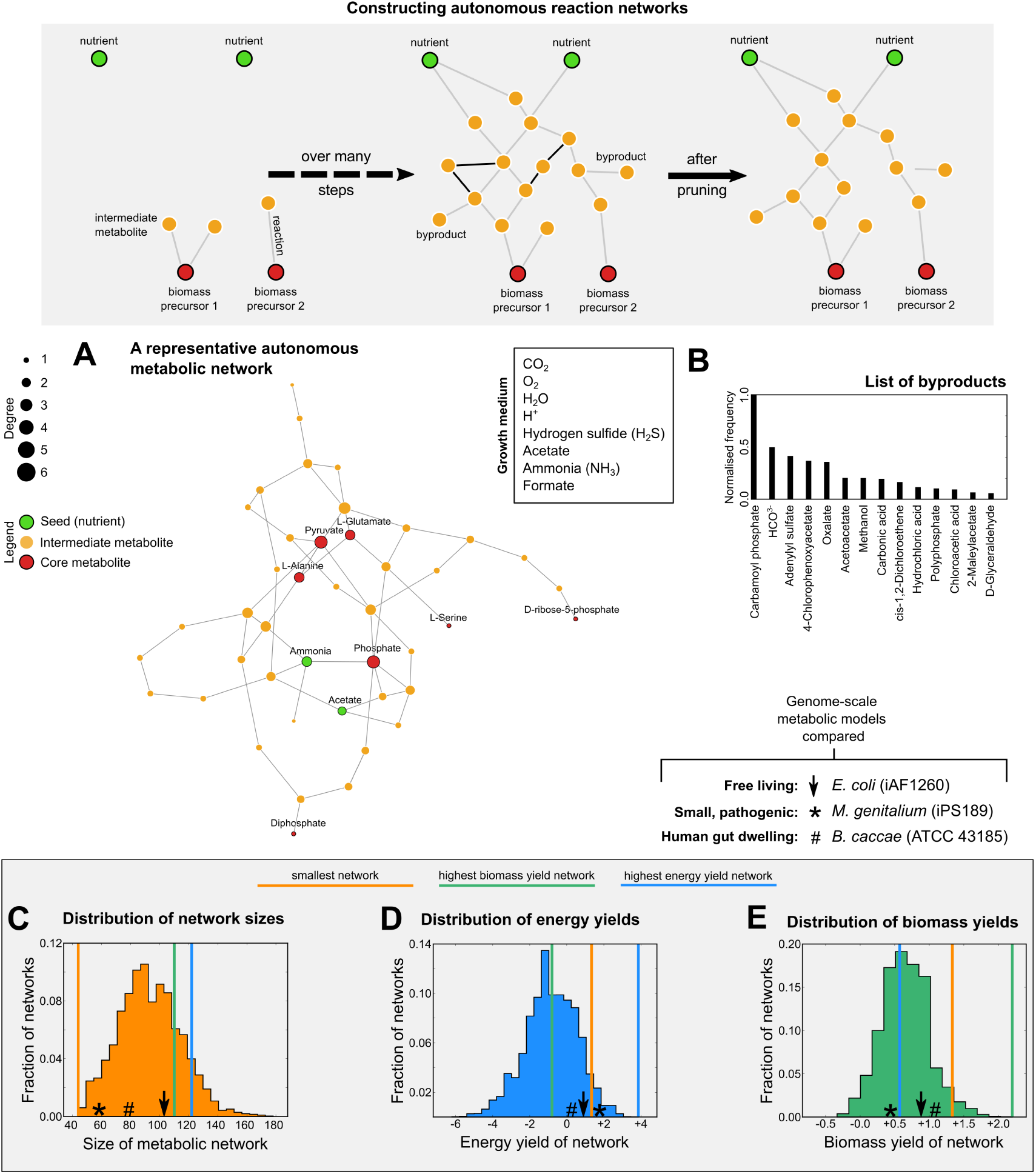
Algorithmically generated autonomous metabolic networks. **(A)** Schematic representation of the algorithm that assembles an autonomous metabolic network (for a detailed description see Methods). **(B)** Network representation of the smallest (having the least number of reactions) metabolic network constructed to be viable on a nutrient medium with acetate and ammonia as the sources of carbon and nitrogen, along with the currency molecules listed in the panel. Each node represents a metabolite, with green nodes representing the nutrients in the growth medium, red nodes the essential biomass precursors, and yellow nodes the intermediate metabolites. For visual clarity, the currency molecules are omitted. The size of each node represents its degree in the network. Links indicate the existence of a reaction that connects the corresponding metabolites, with the arrow pointing from reactant to product. The network shown has 44 reactions (see Table S1). **(C)** List of all byproducts (non-precursor metabolites that are produced but not consumed in a given metabolic network), along with the frequency of their occurrence amongst the 19,543 networks we constructed. **(D, E, F)** Distribution of the sizes, energy yields, and biomass yields (see text for definitions), respectively, amongst the 19,543 unique autonomous networks constructed for this nutrient medium. The networks with lowest size, highest energy yield and highest biomass yields are indicated by thick vertical lines in orange, blue and green respectively (the networks that are smallest and have highest biomass yield are unique, whereas there are 11 networks with the maximal energy yield; the blue line in the size panel corresponds to the minimum size of these maximal energy networks, and similarly the blue line in the biomass panel marks the maximum biomass yield for these 19,543 networks). The symbols on the distributions show measurements from three real bacterial metabolisms (see Methods for how we constructed these three metabolic networks).

Of course, while these 19,543 networks can each produce all the biomass precursors, they may do so with varying efficiency. Therefore, we calculated, for each network, its size (number of reactions in the network), its energy yield (ATP and NAD(P)H generated per nutrient molecule), and its biomass yield (the total amount of precursor molecules generated per nutrient molecule). The yield calculations depend of course on how we distribute metabolic fluxes within the networks. The Methods section describes the simple method we chose, based on stoichiometric demand for each metabolite. Briefly, the method assumes that (i) a metabolite that is a reactant in multiple reactions is distributed amongst them in proportion to the stoichiometry with which it is consumed in each reaction, and (ii) the flux of a reaction with multiple reactants is limited by the reactant coming in with the least flux. We elaborate further on this choice in the Discussion section, but we note here that another method, using a network’s net chemical reaction (see Methods), did not qualitatively affect our results (see supplementary figure 3).

Figures 1C–E show these three distributions – size, energy and biomass yield – over the ensemble of 19,543 autonomous networks we constructed. The smallest network, the ones with the highest energy yield and the one with the largest biomass yield are all different networks (solid vertical bars in figure 1C–E). However, statistically there appears to be no obvious correlation between the three metrics over the sampled networks (supplementary figure 1). Thus, we do indeed observe that the many solutions, for catabolizing the given nutrients into the specified set of biomass precursors, vary significantly in the efficiency with which they perform this task. However, even if we examine only those networks with the maximum energy yield we still find that there are multiple (5) solutions. However, the networks with maximum biomass yield, and with the smallest number of reactions, are unique.

**Table 1:** **Set of reactions for the cross-feeding pair in figure 3A, whose merger gives yield 1.5 (per X)**.

| Reaction network 1 | Reaction network 2 |
| --- | --- |
| $X \rightarrow 3 A + B$ | $X \rightarrow A + BAA$ |
| $A + B \rightarrow BA^*$ | $BAA \rightarrow B + AA$ |
| $A \rightarrow A^*$ | $B + A^* \rightarrow BA^*$ |
| $AA \rightarrow AA^*$ | $2 A \rightarrow AA^*$ |
| **Yield: 2 (per X)** | **Yield: 1.5 (per X)** |

As a consistency check, figures 1C–E also show where three bacteria with fully characterized genome-scale metabolisms – one free-living, one pathogenic and one a human gut dweller – lie on these distributions. All three genome-scale metabolic networks, when pruned to be minimal (see Methods for details), satisfy our stringent survival criterion on this nutrient medium, as well as the alternate medium we have examined where the carbon and nitrogen sources are glucose and glutamate.

Finally, figure 1B shows the set of “byproducts” and their distribution across this set of 19,543 unique autonomous metabolic networks. We define byproducts to be non-precursor molecules that are produced in a metabolic network but not consumed by any reaction. We note that this set is quite small, a reflection of the structure of the KEGG universal chemistry. Further, the average number of byproducts produced in a network is also quite small, typically between 2 and 3 per network (*<* 1% of the metabolites in each network). This will be relevant in the next section where we examine pairs of metabolic networks that are capable of feeding off each other via these “costless” byproduct molecules.

### Obligate cross-feeders can outperform even the most productive autonomous networks

We define a pair of networks to be obligately cross-feeding when both networks require at least one of the partner’s byproduct molecules to produce all the precursors, in order to satisfy our viability criterion. In other words, each network of the pair can perform the metabolic task we impose in the presence of its partner, but not on its own.

To generate random cross-feeding pairs, we started with a randomly chosen pair of autonomous networks that we generated previously, and repeatedly added and removed reaction pathways from them. We did this in such a way that the modified networks required at least one metabolite from each other’s byproducts in order to produce one of the precursors (see Methods for details). As with the autonomous networks, these cross-feeding pairs are minimal, in the sense that removing any reaction renders that network unviable. This procedure successfully resulted in cross-feeding pairs in roughly 1 of 10,000 attempts, and out of 19,543 autonomous networks, 11,358 were unable to form a cross-feeding pair. However, from the remaining 8,185 autonomous networks, we generated 5,421 unique obligate cross-feeding pairs of metabolic networks that satisfied our stringent viability criterion together, but not individually; figure 2A shows one example. We found that in 8% of these pairs, both members have higher energy yields than any autonomous network (shaded region in figure 2B), and 6% have higher biomass yields (shaded region in figure 2C). For the rest of this paper, except where explicitly specified, we will focus primarily on the biomass yields – cross-feeding pairs that have higher biomass yields than the best autonomous networks will be termed “outperformers” and the other cross-feeders, “non-outperformers”. We obtain similar results from energy outperformers.

**Figure 2:**
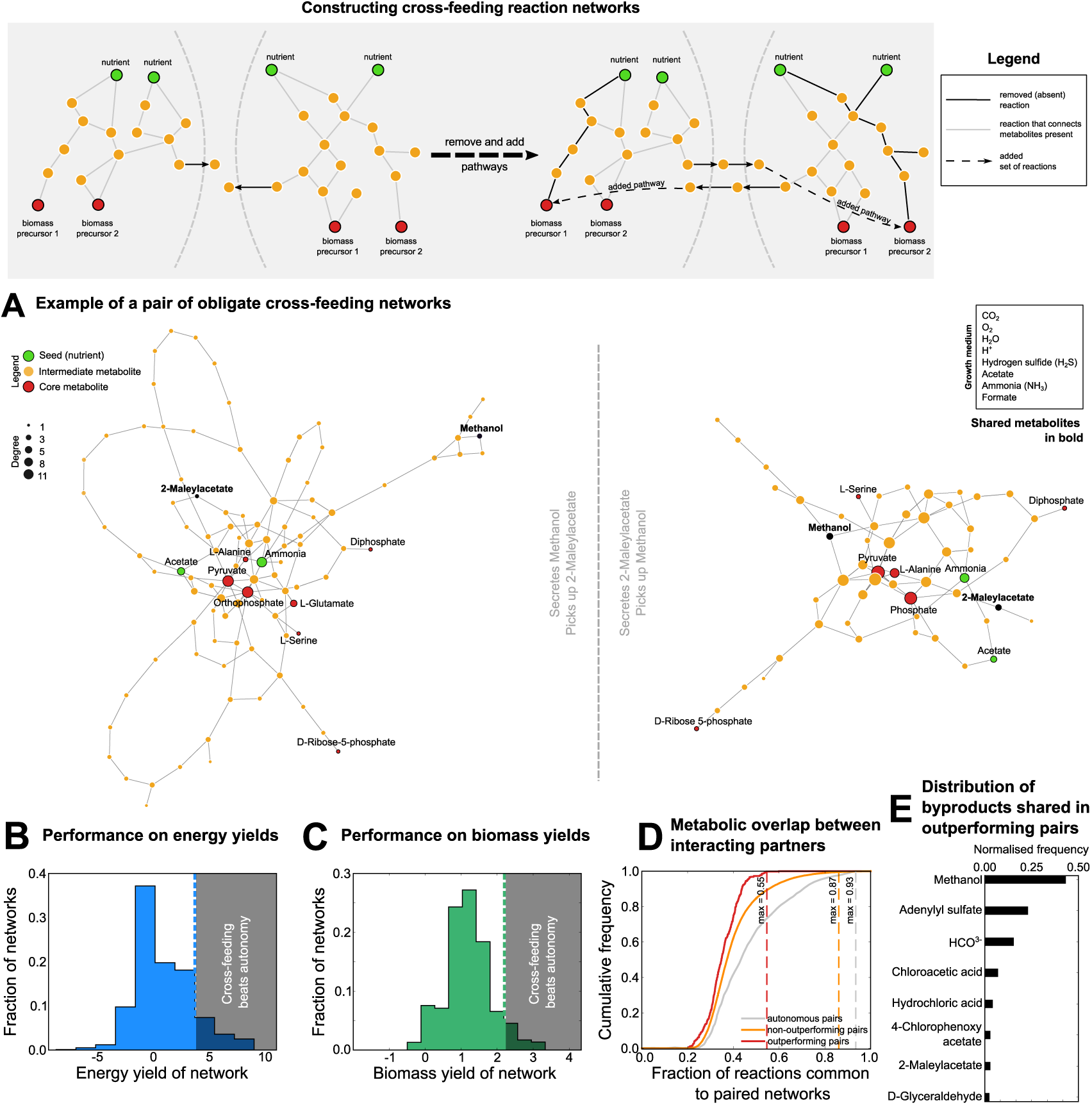
Obligate cross-feeding reaction networks can outperform even the most productive autonomous networks. **(A)** Schematic representation of the algorithm that assembles a pair of obligate cross-feeding metabolic networks (for a detailed description see Methods). **(B)** Network representation of an obligate cross-feeding network pair we constructed. Each node represents a metabolite, with green nodes representing the nutrients in the growth medium, red nodes the essential biomass precursors, and yellow nodes the intermediate metabolites. For visual clarity currency metabolites are omitted. The size of each node represents its degree in the network. Links represent a reaction in the network that connects the corresponding metabolites. The network on the left consists of 89 reactions, and the network on the right has 103 (see Table S2). In this cross-feeding pair, the network on the left secretes methanol and uses 2-Maleylacetate (both in black) from the network on the right (which does the opposite). The distribution of **(C)** energy yields and **(D)** biomass yields of members of the constructed obligate cross-feeding pairs (we plot the lesser of two yields in each pair). The shaded regions marks productivity values beyond those observed from any constructed autonomous network. Cross-feeding pairs that lie in this range we term ‘outperformers’ and the rest ‘non-outperformers’ **(E)** Cumulative distributions of the metabolic overlap (the number of metabolites common to both members in a cross-feeding pair, divided by the total number of metabolites in the union of the two networks) for biomass outperformers (red) and non-outperformers (orange). No outperforming pair has *>* 60% overlap, while non-outperforming pairs can share as many as 87% reactions. **(F)** Distribution of byproducts exchanged between members of biomass outperforming cross-feeding pairs (see supplementary figure 4).

To understand what distinguishes (biomass) outperformers from non-outperformers, we compared several of their structural and biochemical properties, namely: metabolic overlap between partners, number of byproducts exchanged, correlations between yields in a pair, extent of pathway overlap, and responses to various perturbations (see Methods and supplementary figure 5). Strikingly, we found that metabolic overlap – the fraction of metabolites common to both networks in a pair – could help distinguish between outperformers and non-outperformers. Specifically, outperformers typically had intermediate overlap (between 20 and 55%; figure 2D, red), while for non-outperformers, the overlap was typically higher (as much as 87%; figure 2D, orange). Similar numbers were found for energy outperformers (supplementary figure 5). Moreover, we noticed that outperformers typically exchanged only a subset of the byproducts produced by autonomous networks (8, compared to the 14 total byproducts; figure 2E). The set of byproducts exchanged by non-outperformers was larger (12 versus the 14 total). Again, similar observations were made for energy outperformers.

We noticed several additional minor differences between outperformers and non-outperformers. For instance, the yields of the partners in a non-outperforming pair are typically negatively correlated, while those of outperformers are typically uncorrelated (supplementary figure 6). We also found that outperformers and non-outperformers differed in their stability to environmental perturbations, such as nutrient shifts and invasion by other species (supplementary figure 3).

### Choke-points limit key reaction fluxes, and are necessary for outperforming cross-feeders

The existence of cross-feeding pairs that perform better than the best autonomous networks implies that some chemical networks produce a different yield when split into two compartments that interact via a small set of exchanged metabolites. Indeed, we find that the yield distributions obtained by merging each cross-feeding pair into a single network are very similar to the distributions from autonomous networks (see supp supplementary figure 14 for biomass yields). Thus, understanding why the biomass yield of certain pairs of networks is larger than that of their corresponding merged network is key to understanding the existence of outperforming cross-feeding pairs. This is only possible if there are non-linearities in the way metabolic fluxes are distributed. Conversely, if the fluxes in the merged network were a linear superposition of the fluxes of the split networks, then the yields of cross-feeding pairs would be identical to the yields of their merged networks.

We found that the key non-linearity that enables outperforming pairs lies in our stipulation that the flux of a reaction is limited by the reactant which has the minimum flux (see Methods and figure 3). This can result in some key reactions being limited by different reactants when comparing the split and merged networks. We will call such reactions that switch their limiting substrate, “choke-points”. Notice that such a non-linearity can come into play only when there is some intermediate overlap (but not when there is either zero or full overlap) between the metabolites of the cross-feeding pair. If the two networks do not overlap at all, then by definition the yield of the split and merged networks will be the same. Similarly, if there is 100% overlap, i.e., the split networks are identical, then too the yield of the merged network will be the same as the split networks, even if there is a nonlinearity in the distribution of flux. This fits with our observation that outperforming cross-feeding pairs have between 20% and 58% overlap.

**Figure 3:**
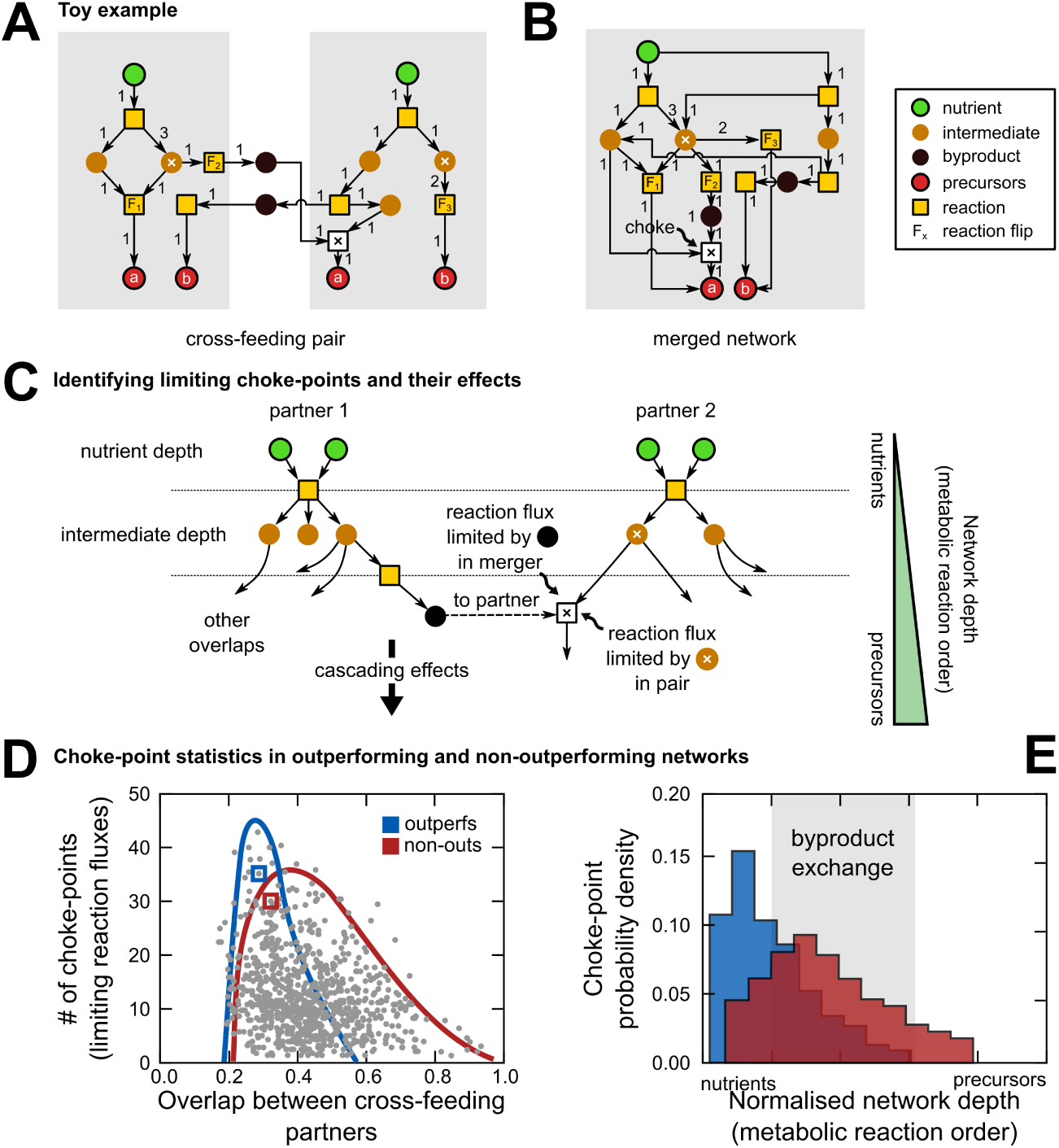
Choke-points limit key reaction fluxes, and are necessary for outperforming cross-feeders. **(A)** A toy example of a cross-feeding pair with a higher biomass yield than its merged network. Both networks use one nutrient (green) to produce two precursors (red, *a* and *b*), via intermediates (yellow). Circles represent metabolites, while squares, reactions. Arrows into a reaction node indicate reactants, and arrows away indicate products of that reaction. A reaction set corresponding to these networks is listed in table 1. Both networks cross-feed off each other’s byproducts (black circles). The total biomass yield of the pair is 2 precursors per nutrient for the network on the left and 1.5 for the network on the right, resulting in a combined yield of 1.75 precursors per nutrient. **(B)** The corresponding autonomous network when the pair in (A) is merged. The metabolite marked with a white cross participates in reactions in both networks (labelled *F*_1_, *F*_2_, and *F*_3_), and is therefore distributed differently in these reactions in the merged network compared to the split pair. Here, this leads to a switch in the limiting substrate for the downstream reaction indicated by the white square with a black cross. This “choke-point”, results in a lower biomass yield for the merged network (1.5 precursors per nutrient). **(C)** Comparison of choke-points in the cross-feeding pairs we constructed and their merged networks. For networks with multiple nutrients, precursors and layers (measured in network depth, i.e., distance from nutrients), it is possible for an overlapping reaction (white cross) to lead to limited flux for a downstream reaction. This “limited substrate switching” can have cascading effects on downstream reactions, resulting in differences in autonomous network performance. **(D)** Scatter plot showing metabolite overlap between both members of 500 randomly chosen cross-feeding pairs versus the number of choke-points (limiting substrate switching events) in them. The blue envelope shows the 95^th^ percentile range of outperformers, and the red, that of non-outperformers. The blue and red squares highlight an example outperformer and non-outperformer, respectively, that have similar overlap and number of choke-points. **(E)** Histograms of the location, in terms of normalized network depth, of identified choke-points in outperformers (blue) and non-outperformers (red). The network depth for a metabolite is the minimum number of reaction steps between the nutrients and that metabolite. The normalized network depth is that number divided by the maximum depth for that network. The gray region shows the range of depths where byproduct exchange occurs in the cross-feeders.

The toy example in Fig 3A illustrates in a simple setting how even a single choke-point can result in split networks producing a different biomass yield than their merged network. Here there is 1 nutrient and 2 biomass precursors (labelled *a* and *b*). Fig 3B shows the merged network (the byproducts, still in black, are now internal, intermediate molecules). The list of chemical reactions corresponding to these networks is listed in table 1. According to our flux distribution scheme, in the merged network the net flux that produces the overlapping metabolite, marked with a white cross, is 4, and this flux is distributed amongst 3 reactions labelled *F*_1_, *F*_2_ and *F*_3_. Following the stoichiometric demand rule, *F*_1_ receives a flux of 1 from metabolite A, *F*_2_ receives a flux of 1, and *F*_3_ receives a flux of 2. From *F*_1_, the metabolite B is split into two downstream reactions. One of these reactions (marked choke in Fig. 3B) is limited by B in the merged network, but the byproduct A* (which is cross-fed) in the split networks. Thus, between the split and merged networks, there is a switch in which reactant limits the choke-point reaction; this switched limitation leads to a difference in how much biomass precursor BA* is produced downstream, which ultimately leads to differences in the yields between the merged and split networks. Since the limiting reactant in the choke reaction has a lower flux in the merged network than the split network, the merged network has a lower biomass yield than the split networks combined (1.5 for the merged, versus 1.75 for the split).

The networks we have generated from KEGG’s universal chemistry typically have many more overlapping metabolites, and are much deeper (median un-normalized network depth, from nutrient to precursor, is 8). Thus, we not only expect many more choke-points to occur in our cross-feeders, but we also expect their effects to cascade much further (figure 3B). For a 1000 cross-feeding pairs (54 outperforming, 946 non-outperforming), randomly chosen from our ensemble of 10,000, we recorded both the number of choke-points (see Methods), as well as their distance from the nutrients (normalized network depth - see Methods) in the merged network. We found that the number of choke-points initially increased with metabolic overlap, reaching a peak at around 40% overlap, and then reduced as the overlap became much larger (figure 3D). Moreover, we found that, on average, not only did outperformers have a lower overlap compared to non-outperformers, their overlapping metabolites were more likely to cause limiting substrate switching (average number of cases of switching being 18 ± 2 versus 12 ± 1; figure 3D). In outperformers, these choke-points are typically located before byproduct exchange (on average, 1–2 layers before; figure 3E), whereas in non-outperformers, these are typically located in the same layers where byproduct exchange occurs (figure 3E, outperformers in blue, non-outperformers in red, byproduct exchange in gray). Together these results show that not only do outperformers typically have more choke-points, but that they also occur closer to the nutrients and away from the precursors, thereby making them capable of increased downstream cascading effects.

To summarize, thus far we have shown that for a pair of cross-feeding networks to produce higher biomass yield than their merged network, they need: (i) a non-linearity in the flux distribution where some reactions switch their limiting substrate, i.e., the existence of choke-points (particularly in the layers between the nutrients and the byproducts); and (ii) an intermediate metabolic overlap, without which this non-linearity cannot manifest itself. These two are *necessary conditions* for such “local” benefit of cross-feeding, where a cross feeding pair does better than their merged network. However, in the previous section we found a stronger “global” benefit, where some cross-feeding pairs performed better than the *best* autonomous networks, not just the network formed by merging them. In fact, we find that these two conditions are also necessary conditions for such global benefit – that is, if a cross-feeding pair does *not* satisfy (i) the probability of it being an outperformer is 3%, and if it does *not* satisfy (ii) the probability is 0%, both of which are close to zero. The few cases that slip through these conditions, especially condition (i), arise because it is possible for the yields of split networks to differ from their merged networks without triggering a switch of a limiting substrate – the same non-linearity that redistributes fluxes is at play in such case, but in a subtler form. However, because the switching of the limiting substrate is what underlies most outperforming cross-feeding pairs, we chose to focus on that more dramatic manifestation of the flux non-linearity.

### Outperforming cross-feeding networks arise from efficient byproduct donors and acceptors

While we find that the existence of choke-points and an intermediate metabolic overlap are necessary conditions for a pair of cross-feeders to outperform, they are not sufficient conditions. That is, if either condition is not satisfied then the pair is almost certainly not an outperformer; but both conditions being satisfied in fact predicts outperformance no better than a coin-flip (41% of cross-feeding pairs that satisfy both these conditions are outperformers; supplementary figure 8). This is not surprising, because while non-linearities may make a split network produce a different yield from its merged network, it could as well perform *worse* rather than better than the merged network. We now explore additional properties that can predict whether the effect of such non-linearities is in favour of cross-feeding or against.

Recall that our algorithm for constructing cross-feeders performs a two-step modification on a pair of autonomous networks (see figure 4A–B for an illustrative example). First, it removes a set of reactions (a metabolic pathway) that produces a specific precursor (figure 4A, shaded region). Then, it adds a compensatory pathway, which uses an external byproduct (figure 4B, shaded region). Pathway removal, the first step, removes branch points at certain nodes (branch partition in figure 4A). This provides higher substrate amounts to the remaining reactions, which increases their flux, and results in higher product amounts (partition relieved in figure 4B). One effect of this is to change the amount (or yield) of the byproducts produced by the modified network (from *Y*_*byp*_ to 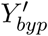, figure 4A). For a good donor, this change must be positive, since it means a higher amount of byproduct produced for its partner. Therefore, to measure the donor quality of a particular autonomous network, we calculated the highest change in byproduct yield that could result from the removal of one its pathways, max 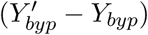 (see Methods).

**Figure 4:**
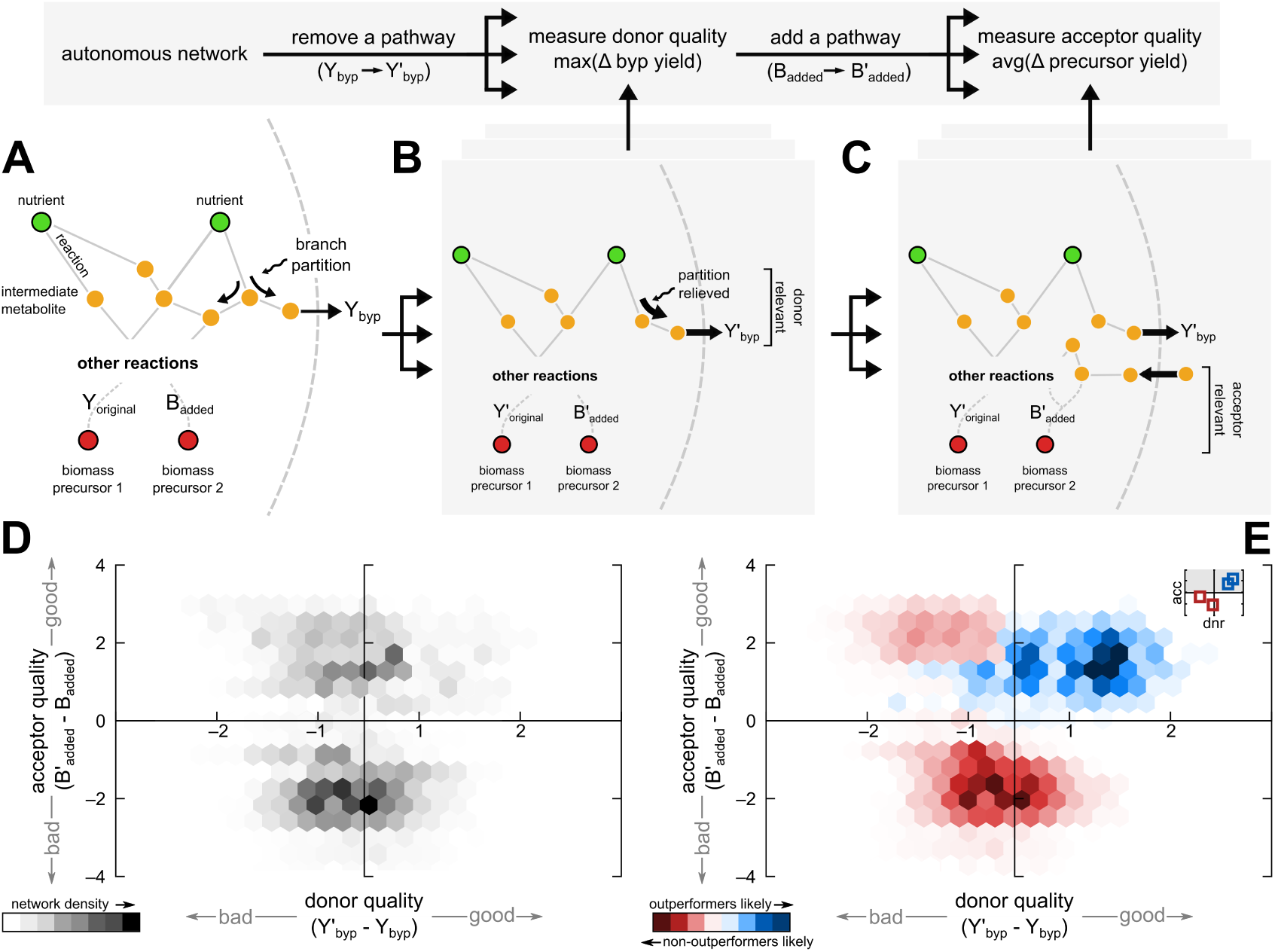
Donor and acceptor qualities of autonomous networks. **(A)** An illustrative example of an autonomous network, that converts nutrients (green) to biomass precursors (red), via intermediates (yellow). Undirected links indicate a reaction that involves two molecules. Intermediate metabolites which are substrates in multiple simultaneous reactions are partitioned by stoichiometric demand. Solid arrows indicate specific reaction fluxes, whose thickness scales with flux amount. Substrate inflow to a reaction determines corresponding product amounts, or yields. Three such yields are shown: one for the generated byproduct, *Y*_*byp*_, and one each for the precursors, namely *Y*_*original*_ and *B*_*added*_. **(B)** A corresponding cross-feeder of the network in figure 4A, through: (1) pathway removal, which relieves a previous branch partition; and (2) pathway addition, which allows external byproduct utilization. (1) changes byproduct yield to 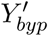, which indicates its strength as a donor. (2) changes its precursor yield to 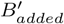, which indicates its strength as an acceptor. **(D)** The donor and acceptor qualities, 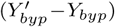 and 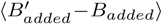 respectively, for all 100,000 autonomous networks generated from KEGG. Networks with similar donor and acceptor qualities are hex-binned, and coloured by their number density. **(E)** The donor and acceptor qualities, 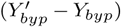 and 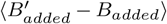 respectively, for 8,185 autonomous networks generated from KEGG, which could be converted to cross-feeders. Networks with similar donor and acceptor qualities are binned together in hexagonal bins, and coloured by their likelihood to become outperformers (blue) or non-outperformers (red).

After the pathway removal step, one of the precursors cannot be produced by the modified network. In order to compensate for this, when we construct obligate cross-feeding pairs we add another pathway that uses an external byproduct to produce this precursor. The reactions in this added pathway yield a certain amount of the missing precursor, say 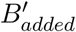. For a good acceptor, such a pathway addition must satisfy two conditions: (1) the change in the amount of precursor produced must be positive; and (2) the added pathway must not compromise the reaction fluxes of the rest of the network, or do so as little as possible. Therefore, to estimate the acceptor quality of a particular autonomous network, we calculated the highest change in its total precursor yield, 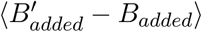, averaged over all possible input byproducts as well as output precursors (see Methods). For each network, the input byproducts considered were the full set of 14 byproducts found in the cross-feeders we generated.

We calculated a donor and acceptor quality for each of the 19,543 unique autonomous networks that we generated. We found a clear bimodality in acceptor quality with the trough between the two peaks being centered at zero (see Fig 4D). In contrast, while the distribution of donor quality spans a broad range from negative to positive numbers, there is no natural separating line between good and bad donors. Nevertheless, we will define “good” acceptors and donors to be those with acceptor, or donor, quality greater than zero. Fig 4E shows the same plot of acceptor vs donor quality for a subset of 8,185 autonomous networks whose modification yielded cross-feeding networks, coloured according to their ability to produce outperformers (blue indicates a high probability, and red a low). From this plot it is clear that outperforming cross-feeders arise largely from autonomous networks that are both good donors *and* good acceptors. Only 4% of networks that do *not* satisfy this condition are outperforming cross-feeders. Therefore, despite being a rather weak requirement (acceptor and donor qualities just need to be positive, not particularly large), this still functions as a necessary condition that we can add to the two others from the previous section. In addition, the acceptor quality of an autonomous network was somewhat correlated with the quantitative yield gain for cross-feeding pairs it generated (see supplementary figure 6).

### Conclusion: necessary and sufficient conditions for a cross-feeding pair to outperform autonomous networks

We can succinctly summarize our results as follows:

1. The following three conditions are each *necessary* conditions for outperforming cross-feeding: (i) metabolic overlap must lie between 20% and 60%; (ii) choke-points must exist; (iii) the autonomous networks from which cross feeders are generated must be good donors and good acceptors. That is, if any one of these conditions is *not* satisfied, the cross-feeding pair is very unlikely to outperform the best autonomous network (top three bars of supplementary figure 7).
2. Together the three conditions are an *almost sufficient* condition for outperforming cross-feeding. More precisely, approx 80% of outperforming cross-feeders satisfy all three conditions (bottom-most bar of supplementary figure 7).

Note that condition (iii) is a relatively weak constraint – it only demands that donor and acceptor quality is greater than *zero*, not that they need to be large.

Satisfying only one or two of these properties is *not sufficient* – supplementary figure 7 shows that satisfying combinations of two of these conditions, but not all, predicts outperformers at best a little better than a coin flip. For a specific example, see figures 3D–E. Here, the outperformer marked by a blue square and the non-outperformer marked by the red squares have similar overlap and number of choke-points (figure 3D). However, their individual source autonomous networks occur in opposite quadrants of the donor-acceptor plot (figure 3E, inset).

The 20% of networks that slip through these three combined conditions consist mainly (bottom-most bar in supplementary figure 7) of outperforming cross-feeders where the donor has negative quality and a few where the non-linearity manifests itself in a more subtle manner, rather than in a switching of the limiting substrate of some key reactions. More complex versions of conditions (ii) and (iii) could easily be constructed that take care of this and account for almost all the outperforming cross-feeders. For instance, we can increase the percentage of predicted ourperformers to 86% if we modify condition (ii) to read: choke-points must exist and *>* 50% of them should lie in the network layers between the nutrients and the byproducts. However, this loses the intuitive simplicity of the original three conditions.

## Discussion

Here, we have explained, using realistic metabolic networks, how metabolically interdependent (cross-feeding) microbes may outperform even the best autonomous microbes at the task of building a set of biomass precursors in a given nutrient environment. This provides a theoretical basis for the empirical observation that pairs of microbes that cross-feed often have a higher growth rate than autonomous microbes [5, 9, 34]. The networks we construct, while randomly assembled, are built out of real chemical reactions. Thus, the underlying chemical basis for metabolic interdependence is clear and, for example, the small sets of metabolites that we find mediate cross-feeding consist of real molecules whose existence in soil, gut or marine ecosystems can be tested. Previous studies have implicated fermentation end-products such as acids, alcohols and acetates, as exchanged metabolites during diffusion-dependent cross-feeding [35, 36, 37]. This is consistent with the metabolites exchanged by the outperforming cross-feeders in our study.

We argue that enhanced biomass or energy yield of certain cross-feeding networks necessarily requires some non-linearity in the way metabolic fluxes flow through the network from nutrients to biomass precursors. In the simple scheme we use to illustrate this, the only non-linearity lies in the flux of a reaction being limited by the reactant coming in with the least flux. This non-linearity would also exist in more complex flux distribution schemes (e.g., those based on flux balance optimisation, or on an explicit regulatory network). The controlling regulatory network may of course introduce other non-linearities, but this one, we have shown, is sufficient to enable beneficial cross-feeding.

We found that for cross-feeding pairs to outperform autonomous networks, three conditions must hold simultaneously: (i) the cross feeding pair must not have too much or too little overlap, (ii) some key reactions must switch their limiting substrate in the split networks compared to the merged network, and (iii) the two cross-feeding networks must be good donors and acceptors of byproduct metabolites. We believe the three conditions we have uncovered are likely to be independent of the particular choice we made of the rules for distributing the metabolic flux in these networks. The reason is that they lay out very general *phenomenological* criteria. The precise form of the non-linearity, which particular reactions switch, and the specific values of the donor and acceptor qualities will change if we use a different flux distribution scheme, but the three criteria will still hold. If the first two conditions do not hold, i.e., if a non-linearity is not triggered, or if the overlap in networks is zero or maximal, then the flux distribution in the merged network will be a linear superposition of the fluxes in the split networks and no benefit can accrue. The donor or acceptor quality quantifies the ability of metabolic networks to benefit from secreting a metabolite they don’t need and from using a metabolite they don’t produce. Thus, the third condition simply encodes the very general idea that the non-linearity must act in favour of, and not against, the split networks. The donor and acceptor quality of a network depends on chemical and thermodynamic properties of the set of reactions in the network. It would be of interest in future work to elaborate precisely what chemical and thermodynamic conditions lead to networks being good donors and good acceptors. We also note that the donor and acceptor quality as we have defined it is in principle measurable empirically, albeit requiring some possibly tedious manipulation of the metabolic pathways of a microbe. Comparing such measurements with theoretical predictions would allow us to understand better how metabolic fluxes are actually distributed by regulatory networks *in vivo*.

The biological consequences of our results are several. First, the existence of outperforming cross-feeders suggests that once such cross-feeders arise in a microbial community they will likely do well, assuming that increased energy or biomass yields (as we have defined them) would correlate with increased growth rate or ability to compete with other strains. It would be useful to have a clean way to quantitatively predict the growth rate of a cross-feeding pair in laboratory conditions from the yield values. But in the absence of additional data, how to do this is not obvious. The growth rate may depend on some weighted combination of the energy and biomass yields, and these weights in turn may depend on the specific environment in which the strains are growing. We did check that if we define productivity to be a weighted combination of the biomass and energy yields, and the size of the network, we still find many outperforming cross-feeding pairs for a wide range of weight choices (see supplementary figure 4C). One way to calculate growth rates could be to introduce a “biomass” reaction such as the ones used in flux balance optimisation. In addition, to predict actual growth rates it may be necessary to include the cost of secreting metabolites, even if they are byproducts. At the moment we do not include such costs, so the yield benefits we obtain for cross-feeders are best thought of as *upper bounds* to the benefit of cross-feeding – the larger the predicted yields, the higher the costs of secretion and uptake that the species pair can sustain and still achieve a net benefit. A useful way of extending our work would be to allow secretion of not just the byproducts but other intermediate metabolites, and even biomass precursors themselves, thereby asking whether cross-feeding could also benefit from exchange of such “costly” metabolites. Despite these caveats, we believe our results nevertheless establish a strong theoretical basis for understanding when cross-feeding can be beneficial.

An obvious question that arises is whether there exists a plausible evolutionary path whereby an initially homogeneous population, consisting of all cells having the same autonomous metabolic network, could spontaneously split into two sub-populations which form cross-feeding pairs. This would require the existence of a series of evolutionary steps, consisting say of a loss or gain of a metabolic pathway, where at each step the resultant strains would be doing at least as well as the autonomous networks. We do find that certain outperforming cross-feeding pairs exist with sufficient (∼ 70%) overlap, such that the difference between them is no more than one or two metabolic pathways. Therefore, our results show that such an evolutionary path is plausible, but more work would be needed to explicitly demonstrate such “symmetry breaking” [**?**, 9, 38, 39, 40].

On a broader note, metabolic interdependence of the sort we have studied is very closely related to the question of compartmentalization of reaction networks. This is relevant not just in microbial communities but also in a variety of other contexts, such as within eukaryotic cells or in industrial contexts. Many different reasons have been proposed for why compartmentalization might be beneficial – for instance, it can prevent the formation of toxins or waste products, it can prevent runaway polymerization, smaller volumes in smaller compartments can alter the effective reaction rates, it can be used to control timing of reactions more precisely, etc. Our work provides another mechanism, via non-linear redistribution of reaction fluxes, that can operate together with these other possibilities. To understand if such flux redistribution is important in say eukaryotic cells, one could replace the KEGG universal chemistry we use (which includes only reactions found in bacteria) with the chemical reactions that are known to occur in eukaryotic cells. This would demonstrate whether compartmentalizing the reactions that occur, for example, in mitochondria produces benefits similar to what we have shown to be possible for bacterial metabolic networks. Similarly, a number of enzymatic cascade reactions have applications in industry, and compartmentalization has empirically shown some success at increasing yields [41]. Our approach could potentially be used to predict optimal ways to compartmentalize such reaction networks, especially because the regulation of the flux is likely to be better characterized, and even controllable, in these industrial contexts.

## Methods

To build a set of chemical reactions from which we will assemble metabolic networks, we extracted chemical reaction data from the microbial metabolism database in KEGG [42]. During extraction, unless explicitly stated otherwise, we assumed that all reactions were reversible. This resulted in what we call the “KEGG universal chemistry”, with 2,550 reactions and 1,059 metabolites. Each metabolic network we construct from this set required a medium (or environment). For this, we required each medium to contain at least one carbon source, one nitrogen source, and a set of 10 always available “currency” molecules: water, carbon dioxide, oxygen, hydrogen, ATP, ADP, AMP, NAD(P)H and formate. For our choice of carbon and nitrogen sources, we selected two kinds of media: a simple medium (with acetate and ammonia) and a complex medium (with glucose and glutamine). From these provided nutrients we required each metabolic network to produce a set of essential biomass precursors in order to be “viable”. For this, we curated two sets of essential biomass precursors (table 2): (1) 8 metabolites found to be common to 58 diverse microbial metabolic networks (from ref. [43]), and (2) 8 metabolites in our KEGG universal chemistry from the biomass composition of *E. coli* [44]. Both sets contain precursors of proteins (e.g., L-Serine), nucleic acids (e.g., D-Ribose-5-phosphate), and carbohydrates (e.g., pyruvate).

**Table 2:** Set of designated precursor and currency metabolites for the constructed metabolic networks. (top) The set of metabolites common to metabolic network reconstructions from whole genomes of 58 diverse symbiotic bacterial species (adapted from ref. [43]), along with their KEGG IDs. These metabolites are designated as the set of biomass precursors for all microbial genotypes in the *in silico* networks. For a valid metabolic network in the model, its biochemical reactions must produce *all* of these precursors using the nutrients in the specified growth medium. (bottom) The set of earmarked metabolites that are used as a ‘currency’, i.e. metabolites that we assume are always present in the medium.

| Metabolite | Classification | KEGG Compound ID |
| --- | --- | --- |
| Pyruvate | Precursor | C00022 |
| L-Alanine | Precursor | C00041 |
| L-Cysteine | Precursor | C00097 |
| L-Glutamate | Precursor | C00025 |
| L-Serine | Precursor | C00065 |
| D-Ribose-5-phosphate | Precursor | C00117 |
| Diphosphate | Precursor | C00013 |
| Orthophosphate | Precursor | C00009 |
| H<sub>2</sub>O | Currency | C00001 |
| CO<sub>2</sub> | Currency | C00011 |
| O<sub>2</sub> | Currency | C00007 |
| H<sup>+</sup> | Currency | C00080 |
| AMP | Currency | C00020 |
| ADP | Currency | C00008 |
| ATP | Currency | C00002 |
| NAD<sup>+</sup> | Currency | C00003 |
| NADP<sup>+</sup> | Currency | C00003 |
| NADH | Currency | C00004 |
| NADPH | Currency | C00005 |

**Table 3:**
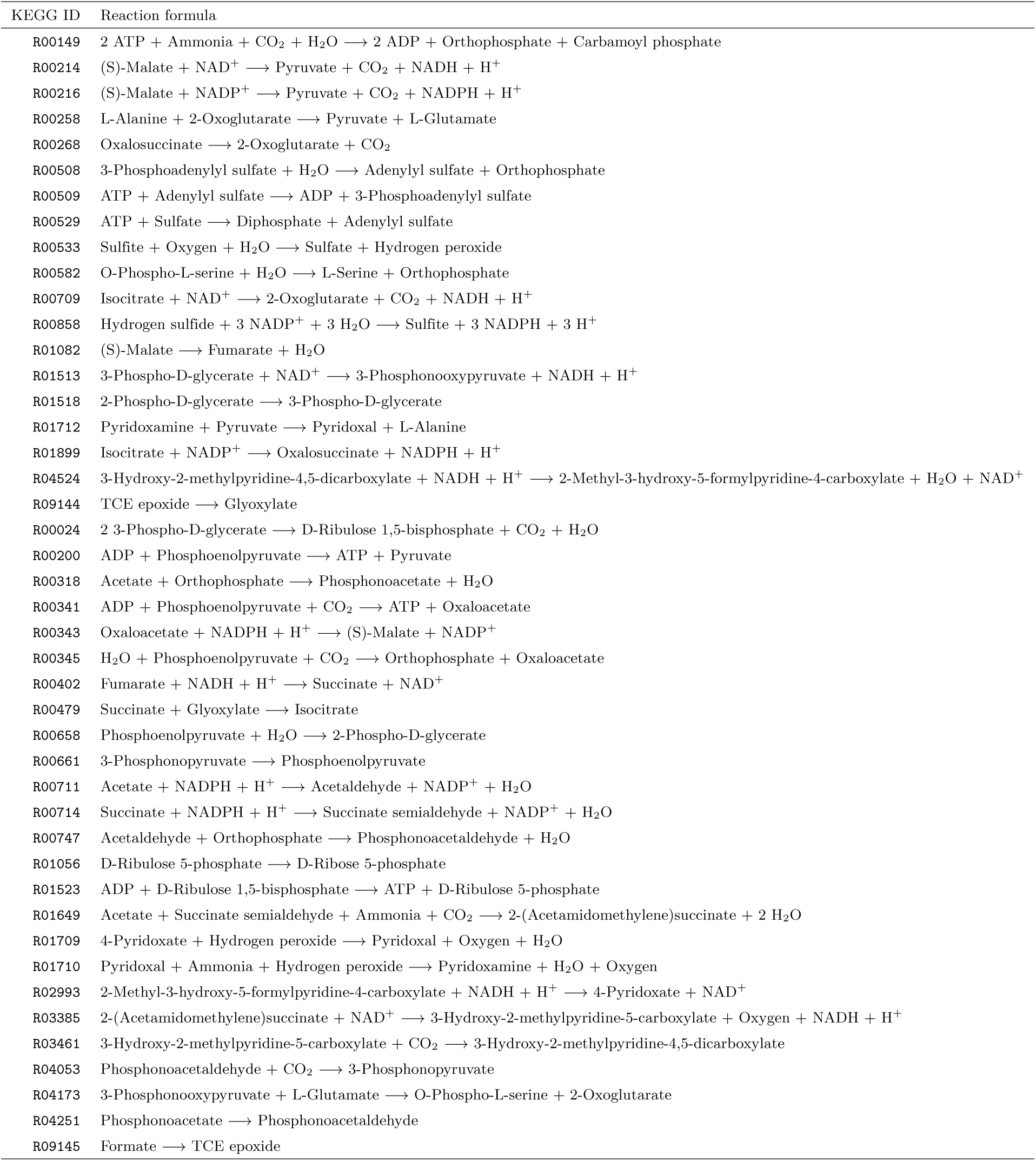
Set of reactions in a sample autonomous network constructed on a sample growth medium. The set of reactions (with their KEGG reaction IDs) in one of the smallest autonomous networks we constructed on a minimal medium with acetate and ammonia. The undirected visual representation of this set is the network in figure 2A. This set has 44 reactions, an energy yield of 1.5 ATP molecules per nutrient and biomass yield of 1.3 precursors per nutrient. Note that in the network constructed, some of these reactions are incorporated in reverse, i.e. the reverse reaction is instead part of the network. In these cases, the correct order is indicated in the column with the reaction formulae.

### Algorithm to construct autonomous networks

For each autonomous metabolic network we generated, we required that it produce *all* 8 precursors (table 2), using only the molecules present in the medium (table 2). We constructed these networks pathway by pathway, i.e., we first construct sets of reactions (“pathways”) which could individually produce each precursor. We constructed “minimal” pathways, that is, pathways where no reactions could be removed without disrupting their ability to generate the target precursor. We did this as follows. We started with a set of reactions denoted *RS* (for reverse scope), which was initially empty. We then queried all reactions in the KEGG universal chemistry which produced the target precursor. We added these reactions to *RS*. Then, we picked all substrates from these newly added reactions, and listed all that were neither nutrients, nor currency molecules. We then queried all reactions in the KEGG universal chemistry that produced each of them. We added these to *RS*. We continued this, step-by-step, until the substrates of all reactions added during a step were present in the medium. We then marked all reactions in *RS* that could be produced using only the molecules in the medium, i.e., the nutrients and currency molecules. We did this by creating a new set of reactions, *FS* (for forward scope), first adding all reactions from *RS* which could be performed using only those substrates present in the medium. At each subsequent step, we added to *F* those reactions from *RS* which could be performed using only nutrients and the metabolites produced by the reactions added to *FS* in the previous step. We continued this till no more reactions from *RS* could be added to *FS*. To make this *FS* minimal, we then pruned it by iteratively removing randomly picked reactions from *FS*, one at a time, such that the remaining reactions could still produce the target precursor starting with only the metabolites in the medium. We then defined the pruned set *FS* as a random minimal pathway which produces a specific precursor. To construct a completely autonomous metabolic network, we repeated this process, generating one pathway per precursor for all 8 precursors. We took the union of the reactions in these pathways, and once again pruned this union as described above. We defined the set of reactions thus obtained as a randomly generated autonomous metabolic network.

### Calculating reaction network yields

We measured the productivity of generated metabolic networks in terms of their energy and biomass yields. A network’s energy yield indicates the number of energy molecules (ATP and NAD(P)H) produced per nutrient molecule consumed, while the biomass yield counts the number of precursors produced per nutrient. We assumed, as per electron transport chemistry, that NAD(P)H molecules provided energy equal to 3 ATP molecules [45], though we verified that changing this ratio does not qualitatively affect our results (supplementary figure 4B). We used two methods to calculate yields for each network, each assuming a different way to distribute the metabolic fluxes: (1) a split-by-demand method, and (2) a net reaction method.

In the split-by-demand method, we assumed that a fixed amount (say, one) of each nutrient molecule was provided to each metabolic network. We first computed which reactions could be performed using just these nutrient and currency molecules. For each such reaction, we used stoichiometric information to calculate which substrate limited the reaction. We assumed that currency molecules never limited a reaction. The amount of products produced by each reaction was determined by its limiting substrate. These products were subsequently used as substrates for the next set of reactions. Here again, the amount of products produced directly influenced which of them limited further reactions. Thus, step-by-step, we calculated the quantities of all molecules generated by the network. At the end of this process, we could sum up and calculate both yields. The network’s energy yield was the net amount of energy molecules generated, divided by the total amount of nutrient molecules provided (2; one each for acetate and ammonia). Similarly, the network’s biomass yield was the net sum of all precursor molecules, divided by the total amount of nutrient molecules provided.

For an example, consider the toy reaction network 1 from figure 3A, which has 4 reactions (Table 1), for which we will calculate the biomass yield. For simplicity, this example has no currency molecules, though when present, these are always assumed to be present and non-limiting. In the first step, only those reactions which consume the nutrient X are possible, which for reaction network 1 is the single reaction X → 3A + B. We assume that X is provided in unit amount, since we normalise yields to the nutrient amounts. Since this reaction uses only one reactant, X, it is the limiting reactant, and therefore the reaction produces 3 units of A and 1 of B. At the next step, two reactions are possible: (1) A + B → BA^*^, and (2) A → A^*^. Both reactions require A, and in equal stoichiometric amounts, because of which we split the 3 available units of A into 1.5 each. Reaction (1) thus has 1.5 units of A and 1 unit of B available, and is limited by B. It therefore produces 1 unit of BA^*^. Reaction (2) produces 1.5 units of A^*^. In this example, the last reaction AA → AA^*^, cannot be completed because the network cannot generate AA itself. When AA is provided from a cross-feeding partner (reaction network 2 in Table 1), this reaction will proceed with a flux determined by the amount of AA provided by the partner. In case no AA is provided, reaction network 1 cannot produce both precursor molecules (BA^*^ and AA^*^), and therefore is not viable, and has no defined yields. If we assume that AA is somehow present, say in unit amount, then the network produces 1 unit of AA^*^. The total yield of this network is 2 (2 units of precursors: 1 of BA^*^ and 1 of AA^*^, for every unit of nutrient X provided).

In the net reaction method, we generated a “net reaction” for each metabolic network. For this, we listed all reactions present in the network once, and summed all the substrates and products on each side. We used this net reaction to calculate the yields of the network. The network’s energy yield was the net amount of energy molecules produced (i.e. the sum of coefficients of ATP and thrice NAD(P)H on the product side) divided by the net amount of nutrients consumed (i.e. the sum of coefficients of the carbon and nitrogen source on the substrate side). Similarly, the network’s biomass yield was the net amount of precursor molecules produced (i.e. the sum of coefficients of all precursors on the product side), divided by the net amount of nutrients consumed (sum of coefficients of all nutrients on the reactant side).

In the example of reaction network 1 in Table 1, the net reaction, obtained by summing over all four constituent reactions, will be: X + 2 A + B + AA → 3 A + B + A^*^ + BA^*^ + AA^*^. The amounts of precursors (BA^*^ and AA^*^) produced are 2 (1 + 1) on the product side, and the amount of nutrient X used on the reactant side is 1. Thus, the total biomass yield of reaction network 1 using this method is 2.

### Algorithm to construct cross-feeding metabolisms

We constructed pairs of obligate cross-feeding networks, where both networks in a pair required at least one byproduct from their partner to be able to produce all precursors. To construct such pairs, we first generated two random autonomous metabolic networks, as described previously. We then picked one of these networks at random, and calculated its byproducts (defined as metabolites that are produced by the network but not used as substrates in any reaction). We then constructed a new pruned pathway which used one of these byproducts as a substrate, and produced one of the 8 biomass precursors. We substituted the corresponding pathway (producing the same precursor) in the second network with this newly generated pathway. The second network now depended on the first. We repeated this pathway substitution procedure, this time generating a pathway using one of the second network’s byproducts as a substrate. When we could perform all these steps successfully in a pair of autonomous networks, we obtained a pair of obligate cross-feeding metabolic networks.

### Comparing with genome-scale metabolic models of three bacteria

To compare the metabolic networks generated by our algorithm with those of real microbes, we extracted 3 genome-scale metabolic models: the free-living *Escherichia coli* [46], the pathogenic *Mycoplasma genitalium*, which has the smallest-known naturally occurring genome [47], and the prevalent human gut resident *Bacteroides caccae* [48]. From each metabolic model, we extracted the EC numbers corresponding to their constituent metabolic reactions, and mapped them against reactions in our KEGG universal chemistry. For *E. coli*, this resulted in 1,339 reactions with 1,039 metabolites. Next, since we assumed reaction reversibility in KEGG’s universal chemistry, we checked if any extracted reactions should be reversed. For this, we replaced the minimum number of reactions with their reverse reactions, such that the resulting reactions could produce all precursors required for growth. Then, to make each reaction set minimal, we pruned this reaction set several times. We pruned in steps; each step, we removed a reaction from the set, and checked if it was still viable given our viability criteria. If it was still viable, we continued the next step with the reduced set of reactions, otherwise we added the reaction back. We continued to prune each reaction set until we could remove no more reactions from it, at the end of which we had a resulting minimal reaction set. From the many (~1,000) times we pruned a reaction set, we obtained a set of corresponding minimal reaction sets, of which we chose the smallest set (or network). We then measured the energy and biomass yields of this smallest corresponding minimal network as described previously (the network for *E. coli* has 104 reactions and 128 metabolites). We used the same procedure to compare two reported cross-fed genome-scale metabolic pairs with our constructed cross-feeding networks as well (figure 1C–E).

### Estimating donor-acceptor characteristics of autonomous networks

To measure the donor quality of an autonomous network, we calculated the maximum difference in byproduct yield upon pathway removal. For this, we first calculated the yield of each byproduct in the network, using the method described previously. We then generated several modified versions of this network, where each version had one pathway removed. These modified networks could not produce a specific precursor. For each such network, we calculated the yields of each of their byproducts. We then calculated the change, or difference, in the yields of byproducts common to both that network and the original autonomous network. We then defined the largest change across all these modified networks as that autonomous network’s donor quality.

To measure the acceptor quality of an autonomous network, we calculated the maximum difference in precursor yield upon pathway addition. For this, we first calculated the yield of each precursor in the network. We then selected 1 byproduct from the 14 listed in figure 1B, and 1 of 8 precursors from table 2. We then used our algorithm to construct several (upto 10, when possible) possible pathways that used the chosen byproduct as a substrate to produce the chosen precursor. We then replaced the pathway that produces this precursor in the autonomous network, with each of our constructed pathways, and measured the change in the yield of the chosen precursor. For this, we assumed that a unit amount of byproduct was available. We recorded the largest of these changes. We then repeated this for each possible byproduct-precursor pair. We defined the average of our recorded yield changes, across different byproduct-precursor possibilities, as the acceptor quality of that autonomous network.

### Calculating predicted and observed yield gains for cross-feeders

For each autonomous network that could be converted to an outperforming cross-feeder, we compared their mean observed yield change with our prediction from their donor-acceptor qualities. For each such autonomous network, we calculated the change in biomass yield, averaged over all outperformers they were part of in our 10,000 generated cross-feeders. We defined this as the average observed yield gain for such a network.

### Detecting limited substrate switching and choke-points while merging cross-feeders

To detect cases of limited substrate switching, we compared cross-feeding pairs with their merged autonomous counterparts, in the following way. For both networks in a given cross-feeding pair, we first recorded which substrates limited each reaction in them, using the split by demand procedure for measuring yields. We then constructed a merged, autonomous version of this pair, by taking the union of reactions in both networks in the pair. We repeated our calculation of limiting substrates in each reaction in this merged network. We then noted in which reactions this limiting substrate were different between the merged network and cross-feeding pair, and defined each such reaction as a case of limited substrate switching. We identified the metabolites that both participated in these reactions and were common to members of the pair, as choke-points.

### Measuring network depth in merged cross-feeders

To measure the location of instances of limited substrate switching, or choke-points, when a cross-feeding pair was merged, we first identified these reactions in the merged network as described previously. Then, for each identified reaction, we measured its depth in the merged metabolic network. For this, we asked at what step this reaction could be performed by the network. Those reactions whose substrates were only the nutrients in the medium and the currency molecules formed the first step. Those reactions whose substrates were either these, or the products of the first step, formed the second step, and so on. At the last step, all precursors would have been produced. We then normalized the position of each reaction by dividing the step at which it could be performed by the total number of steps in the network. We defined this number as the reaction’s depth in the network. To calculate the location of byproduct exchange, we measured the depth (in the merged network) of those reactions that produced and consumed the metabolites that were the exchanged byproducts in the split networks.

## Acknowledgements

We are grateful for many discussions with Szabolcs Semsey and Sergei Maslov that inspired this work. A. G. and S.K. acknowledge funding support from the Department of Atomic Energy, Government of India, under Project Identification No. RTI 4006, and the Simons Foundation (Grant No. 287975). S. K. also acknowledges funding by the Science & Engineering Research Board, Department of Science & Technology, Government of India (Matrics grant MTR/2020/000253). A.G. is supported by the Gordon and Betty Moore Foundation as a Physics of Living Systems Fellow through grant number GBMF4513.

## Figure Supplements

**Figure S1:**
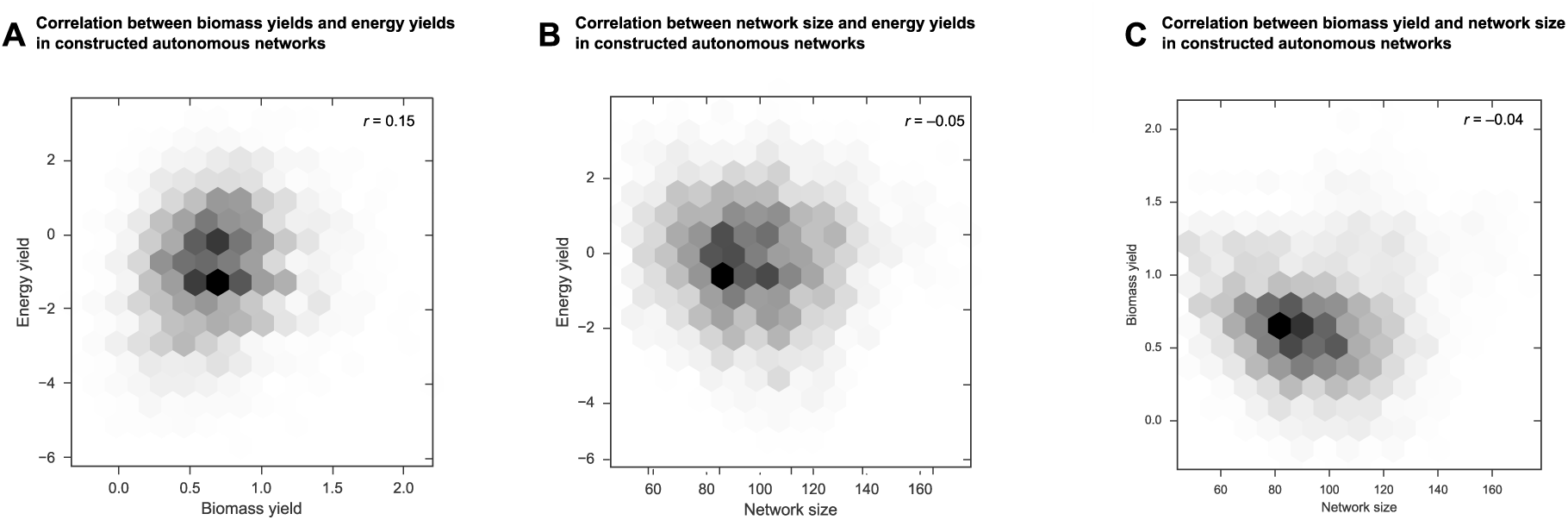
Statistics of sampled energy yields, biomass yields and sizes. Joint density hexagonal plot for the **(A)** energy yields and biomass yields; **(B)** energy yields and network size; and **(C)** biomass yields and network size, for the autonomous networks sampled in a minimal growth medium. The intensity of the color of each hexagon corresponds to the density in that region of the space (the shades are arbitrary since the full density plot is normalized). No two measurements are pairwise correlated in the sampled networks (Pearson’s correlation coefficient, *r* = 0.15, *-*0.05 and *-*0.04, respectively).

**Figure S2:**
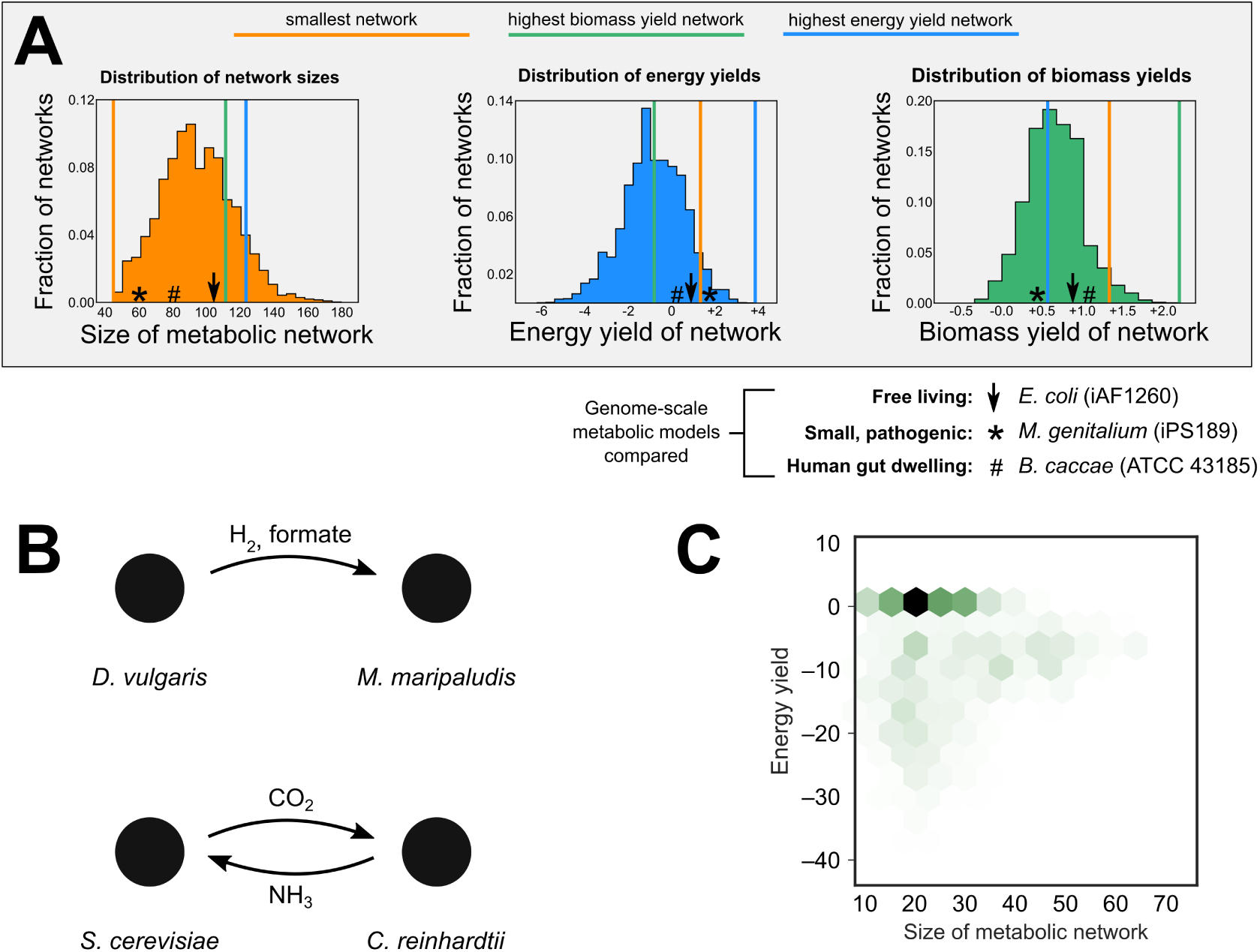
Comparing sampled networks with known genome-scale metabolic reconstructions. **(A)** A comparison of the structure and productivity metrics (as in figure 1D–F) for the sampled autonomous networks in our survey with three genome-scale metabolic models from different lifestyles: the free-living *Escherichia coli* (bold black arrow) [46]; the pathogenic *Mycoplasma genitalium* (asterisk) which has the smallest-known naturally occurring genome [47]; and the prevalent human gut resident *Bacteroides caccae* (hash) [48]. All corresponding reaction networks are viable under our survival criteria. **(B)** We test two reported metabolically interacting strains and their genome-scale metabolic models and find that our model correctly predicts the metabolic interaction underlying them: [top] *M. maripaludis* requires H_2_ and formate from *D. vulgaris* (as in [49]) for pyruvate production, and [bottom] *S. cerevisiae* and *C. reinhardtii* cross-feed CO_2_ and NH_3_ (as in [50]) for pyruvate and amino acid production respectively. **(C)** Joint density hexagonal plot for the energy yields and sizes for metabolisms constructed by randomizing the overall prokaryotic reaction network in KEGG. The networks are much smaller and typically have negative energy yields (which renders them biochemically not viable).

**Figure S3:**
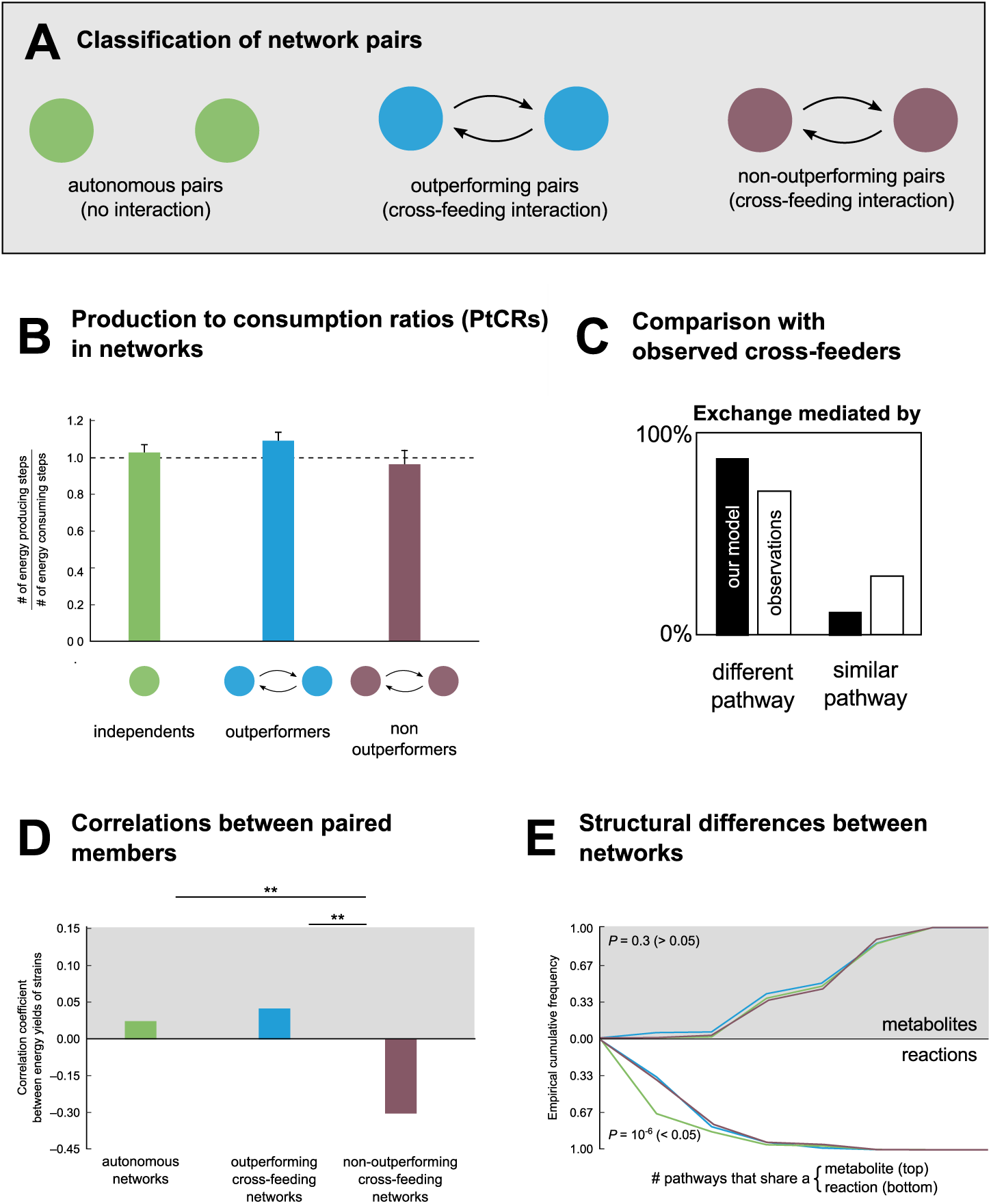
Comparing independent and interdependent networks. **(A)** We compare the networks sampled from algorithmic construction. We classify these as independent (autonomous pairs; green) and interdependent (further split into outperforming cross-fed pairs in blue and non-outperformers in violet) network pairs. **(B)** The production-to-consumption ratios (PtCR; the ratio of the number of energy producing (net ATP gain) to energy consuming (net ATP loss) reactions) for the reaction networks. **(C)** Comparison between constructed cross-feeders with engineered cross-feeders in [51]: the networks from our survey use a similar fraction of pathways that produce different precursors in partners versus the same precursor. **(D)** Correlation between the energy yields of the members in an independent or interdependent pair: we find (as observed in [51]) that non-outperformers usually show a negative correlation. **(E)** Cumulative distributions of the number of precursor molecules that sampled networks whose production requires a common (randomly chosen) [top] reactant and [bottom] reaction. The distributions are plotted for 1,000 randomly chosen possible reactants and reactions respectively, for each category of networks. We compare the cumulative distributions pairwise using a two-sample Kolmogorov-Smirnov test with *P* -value threshold 0.05. We find that for reactants, they do not differ significantly (*P* = 0.3 *>* 0.05) while for reactions, the outperforming cross-feeders are different when compared to both non-outperforming and autonomous networks.

**Figure S4:**
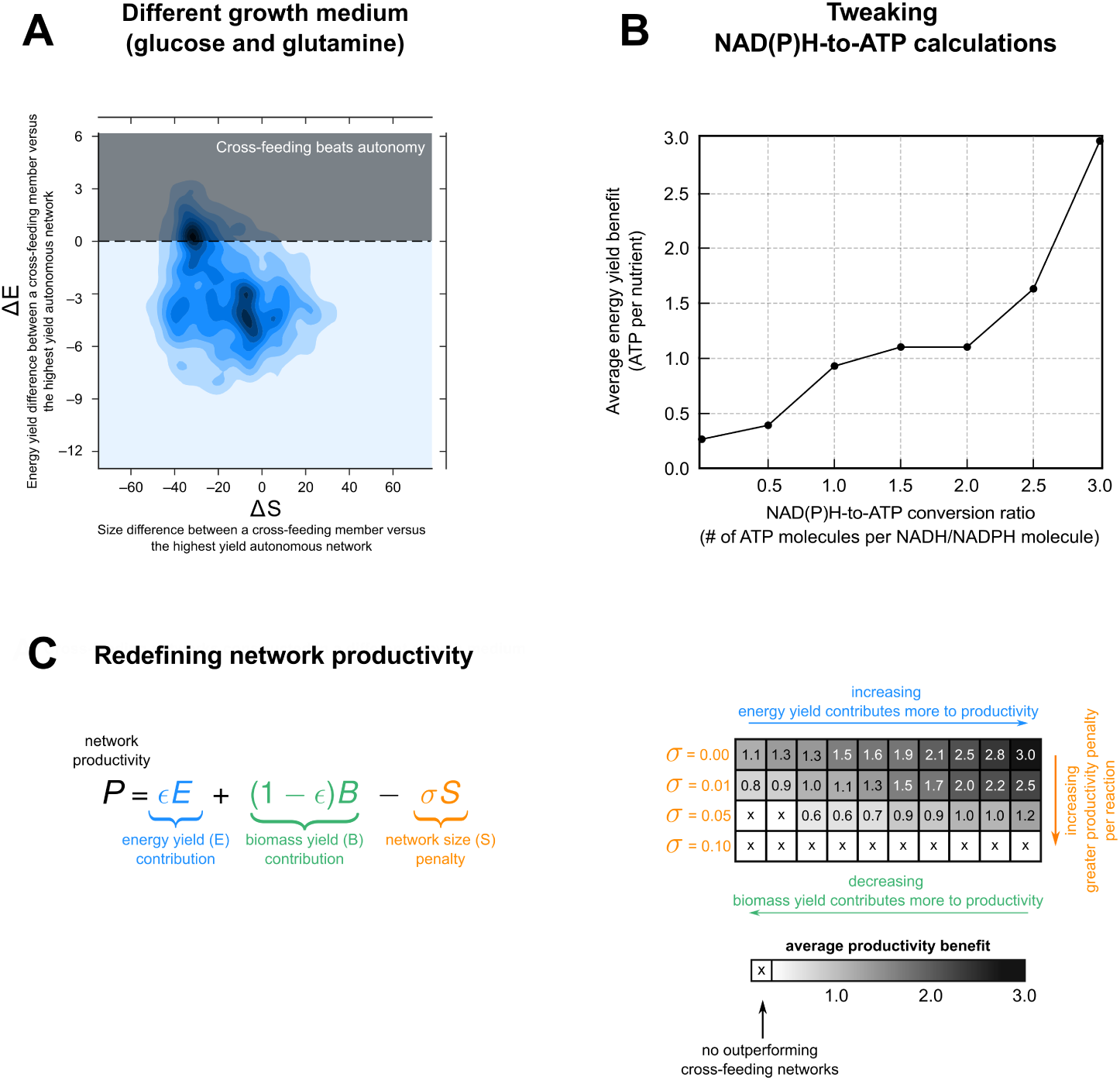
Checking result robustness. **(A)** A joint density plot of Δ*E* on the *y*-axis versus Δ*S* on the *x*-axis for the cross-feeders constructed with a different growth medium (with glucose as the chief carbon source and glutamine the chief nitrogen source). The Δ represents the difference in energy yields (or size) of the cross-feeding versus the highest energy yield autonomous network. Outperforming cross-feeders (Δ*E >* 0) fall above the *x*-axis (shaded region). **(B)** To calculate energy yields for constructed reaction networks, we assume an NAD(P)H-to-ATP conversion ratio of 3. However, there are likely to be cases in which these chains are not as efficient. In these cases, the said ratios will be lower. To test if our central result critically depends on the particular conversion ratio we consider, we repeat our survey with different ratios. Here we show the average energy yield benefit for outperforming pairs (nonzero if it is possible to construct them) as a function of different exchange ratios. **(C)** We redefine network productivity *P* in several ways (via the relation on the left — connecting the energy yield *E*, biomass yield *B* and network size *S* via two parameters *ε* and *σ*) to check if it crucially influences our central result (see *Materials and methods: Result checking with different productivity measurements*). Here we show the average productivity benefit o 0f the sampled cross-feeding networks that outperform the strongest autonomous networks in each parameter set. Each row of the grid represents a fixed value of *σ*, and along a row, each column represents a particular value of *ε*, increasing from 0 to 1 moving along the right. The number in each cell is the average productivity benefit of a cross-feeder. Where it is not possible to construct such a network, we mark the cell with a cross.

**Figure S5:**
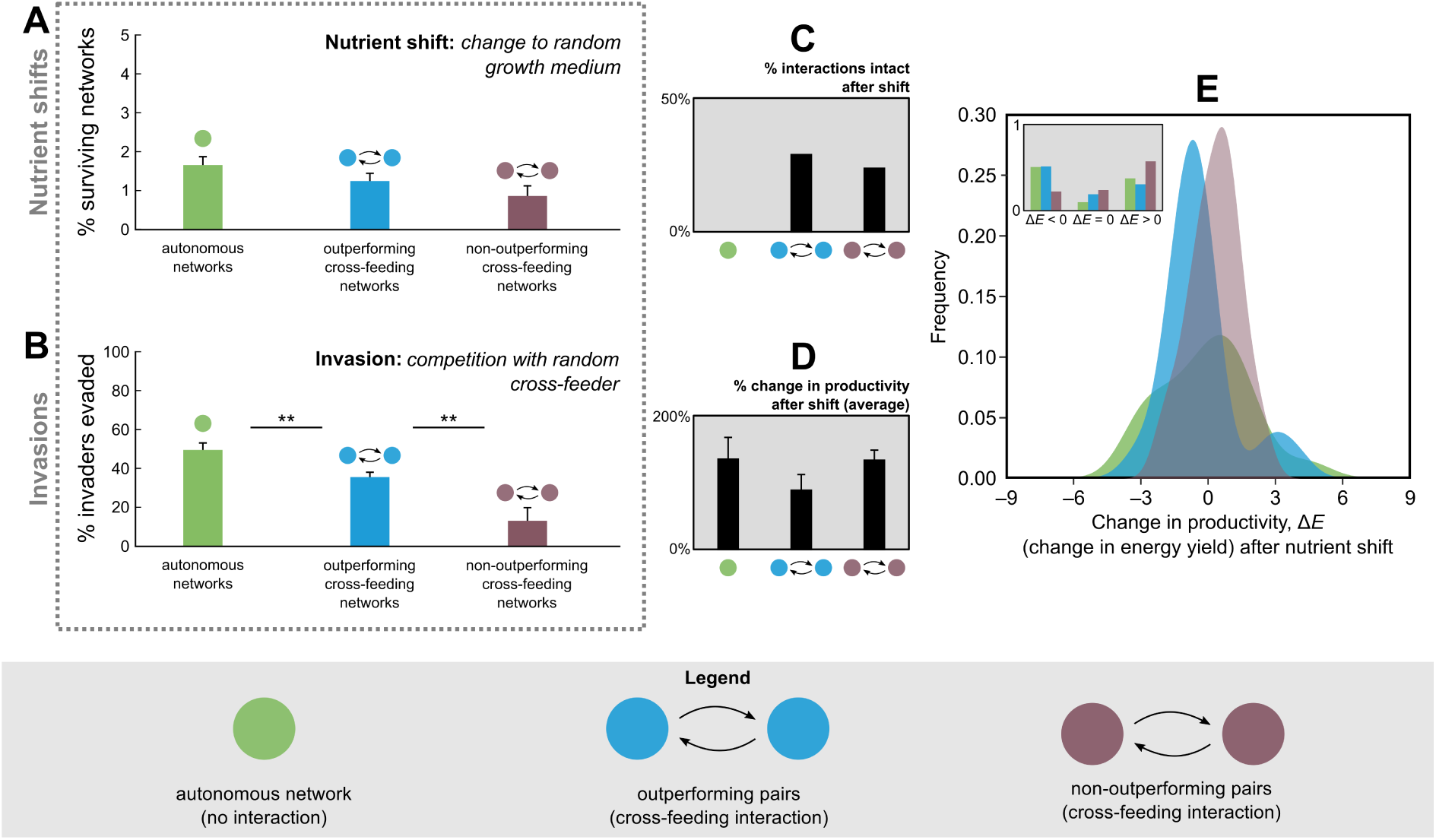
Stability analyses against environmental perturbations for sampled networks. We check how often our sampled reaction networks (autonomous in green, outperforming cross-feeders in blue and non-outperformers in violet) “survive” (i.e. can produce all biomass precursors) under two plausible environmental perturbations: **(A)** nutrient shifts, where we change one of the nutrients in the medium to an alternate reactant in the perturbed reaction network; and **(B)** invasions, where we pit the networks against a randomly chosen outperforming cross-feeder (vis-a-vis their energy yields). Outperformers survive less often than autonomous networks (*P <* 0.01; Student’s *t*-test), but not by a lot (37% as opposed to 50%). Of the surviving networks in (A), we also measure **(C)** how often the cross-feeding interaction is retained, as well as the **(D)** average relative change and **(E)** absolute changes in productivity (energy yield, Δ*E*) after the nutrient shift. The inset classifies the Δ*E*s into negative, positive or no change.

**Figure S6:**
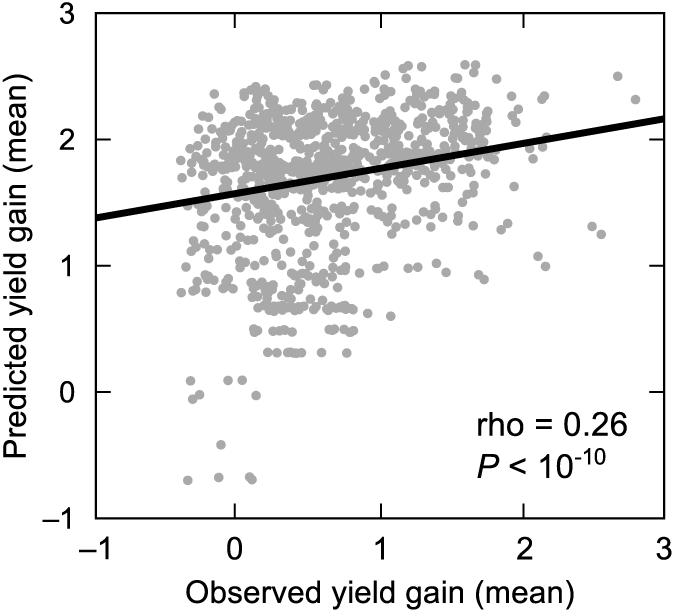
Comparing predicted and observed yield gains using donor-acceptor characteristics. Scatter plot of predicted and observed yield gains for 939 autonomous networks that led to outperformers, with each point representing one network. The predicted yield gain for each network was calculated using its donor-acceptor characteristics. The solid line represents a linear regression (Spearman’s rho = 0.26; *P* value *<* 10^−10^).

**Figure S7:**
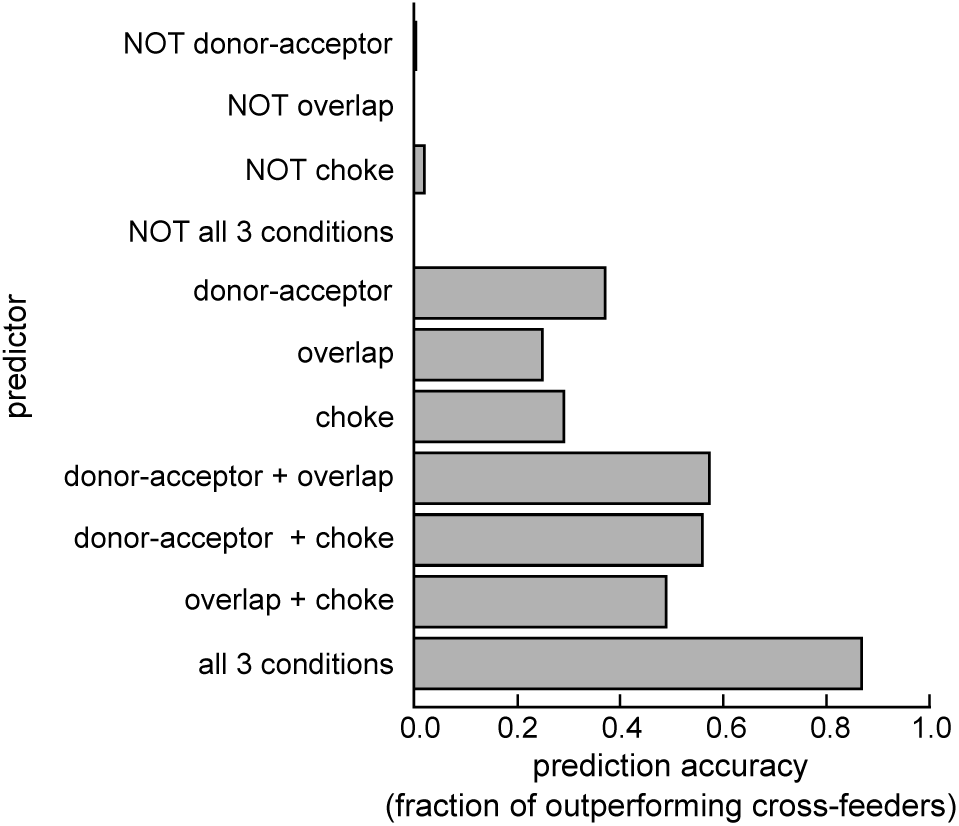
Predicted conditions under which cross-feeders can outperform. Prediction accuracy for conditions that we found were associated with outperforming cross-feeders in our model. For each condition (or predictor), we asked how many of the 10,000 cross-feeding pairs that satisfied it, were outperformers. We tested each condition separately, in pairs, and altogether. The “donor-acceptor” condition measures which networks in the pair arose from autonomous networks that were both in the positive quadrant of the donor-acceptor plot (figure 4C). The “overlap” condition measures whether both networks have between 20 - 60% metabolic overlap (as in figure 3D). The “choke” condition measures which networks in the pair have *>* 50% of their choke-points located prior to byproduct exchange (as in figure 3A).

**Figure S8:**
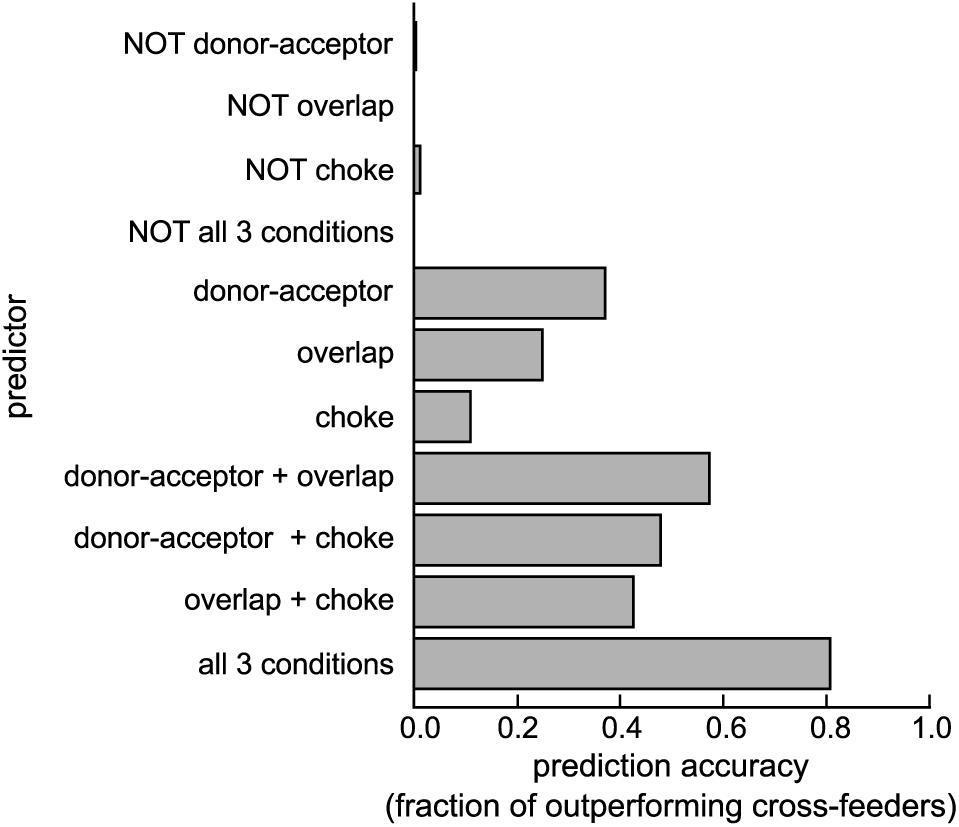
Predicted conditions under which cross-feeders can outperform (with a local choke definition) Prediction accuracy for conditions that we found were associated with outperforming cross-feeders in our model. For each condition (or predictor), we asked how many of the 10,000 cross-feeding pairs that satisfied it, were outperformers. We tested each condition separately, in pairs, and altogether. The “donor-acceptor” condition measures which networks in the pair arose from autonomous networks that were both in the positive quadrant of the donor-acceptor plot (figure 4C). The “overlap” condition measures whether both networks have between 20−60% metabolic overlap (as in figure 5D). The “choke” condition measures whether in a pair, all reactions that used the exchanged byproducts are choke-points, i.e., whether in a pair, all reactions with exchanged byproducts are reactants, switch their limiting reactant when both networks in the pair are merged (as in figure 5A).

**Figure S9:**
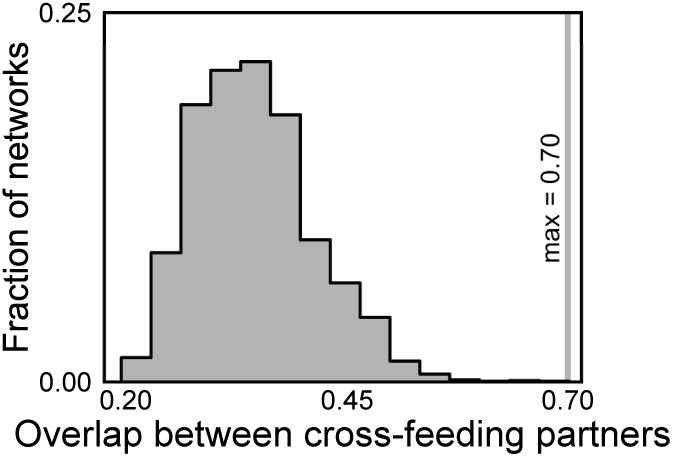
Cross-feeders with better yields than their autonomous counterparts can have higher overlaps. Distribution of metabolic overlap, defined as the fraction of reactions common to both members of a cross-feeding pair, when the yields of both members of a pair were higher their corresponding autonomous networks. This was true for 2,325 of the 10,000 cross-feeding pairs generated. The solid grey line represents the highest such overlap, at 70%.

**Figure S10:**
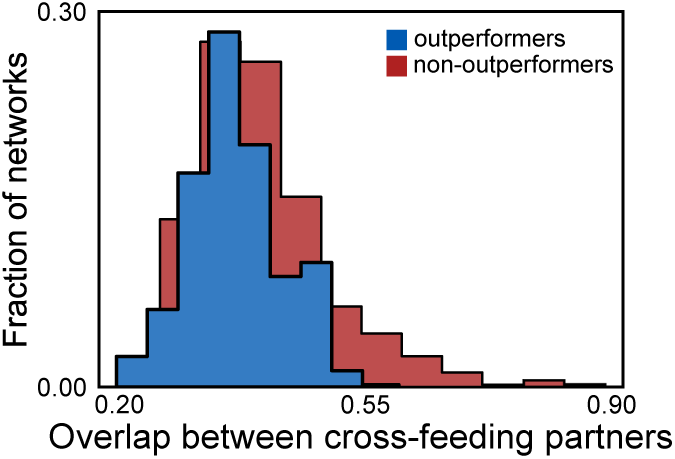
Differences between overlap distributions of outperformers and non-outperformers. Distribution of metabolic overlap, defined as the fraction of reactions common to both members of a cross-feeding pair, of both members of a cross-feeding pair, both outperformers (blue) and non-outperformers (red).

**Figure S11:**
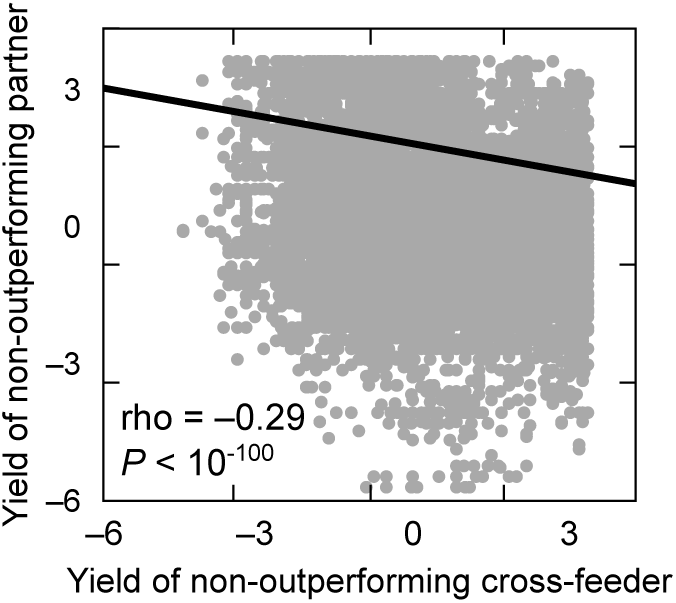
Yields of non-outperforming cross-feeders are negatively correlated. Scatter plot showing the energy yields of both members of all non-outperforming cross-feeding pairs (one member on the

**Figure S12:**
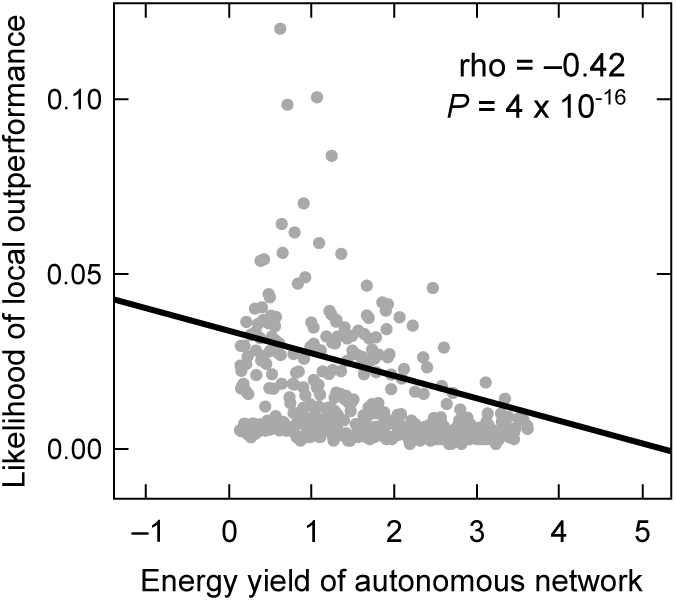
Yields of autonomous networks is negatively correlated with their likelihood of outperforming as a cross-feeder. Scatter plot showing the yield of all 1,000 randomly chosen autonomous networks that could be successfully converted to cross-feeders on the *x*-axis, and the likelihood that (or fraction of) their corresponding cross-feeders had better yields than the autonomous networks themselves. The solid black line represents a linear regression, and rho the Spearman correlation.

**Figure S13:**
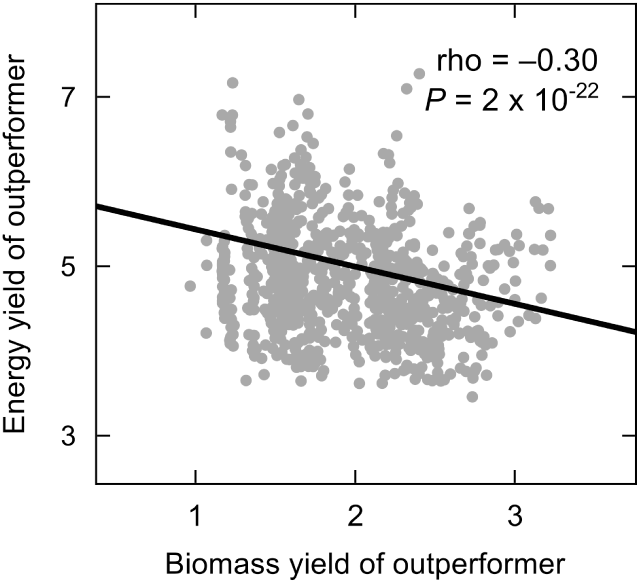
Energy yields and biomass yields of outperforming cross-feeders are negatively correlated. Scatter plot showing the energy yield and biomass yields of all 939 outperforming cross-feeding networks. The solid line represents a linear regression, and rho the Spearman correlation.

**Figure S14:**
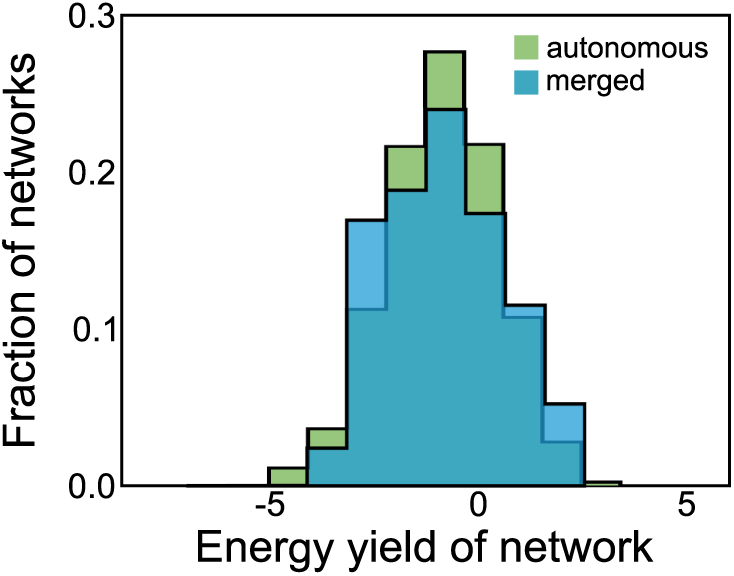
Comparing merged cross-feeders with autonomous networks. Distribution of the energy yields of the 100,000 constructed autonomous networks (green) and the 10,000 merged cross-feeders (blue). When merged, cross-feeder outperformance decreases, and the corresponding networks (also autonomous) do not globally outperform.

